# Engineered Tn5 Variants Enhance CUT&Tag Fidelity for Transcription Factors

**DOI:** 10.64898/2026.10.27.762201

**Authors:** Lola Piper, Emma Baird, Harry J. Ashbaugh, Olivia A. Pentecost, Sara M. Nowinski, Derek H. Janssens

## Abstract

The CUT&Tag method performs exceptionally well for profiling histone modifications, but mapping sequence-specific transcription factors (TFs) is complicated by intrinsic Tn5 activity at accessible DNA, and the high-salt washes used to suppress this activity can destabilize TF-chromatin interactions. We engineered the Tn5 DNA-binding channel to attenuate target-DNA engagement by replacing individual positively charged amino acids with neutral amino acids. We identified a “Goldilocks” variant (K160A) with intermediate attenuation that improved recovery of GATA2 motif-containing sites under low-salt CUT&Tag and KCl-based DynaTag conditions. A second variant (K249A) had improved performance for profiling CTCF over less accessible sites characterized by sharply focused short-fragment enrichment. By comparing the K160A and K249A variants, this TF-CUT&Tag assay provides a physical readout of the local chromatin environment at TF binding sites. Applying K160A TF-CUT&Tag to pluripotency-associated TFs in mouse embryonic stem cells increased recovery across their genomic binding landscapes while preserving factor-specific motif architectures. During skeletal muscle differentiation, TF-CUT&Tag improved recovery of MyoG and Pax7 occupancy relative to standard CUT&Tag, revealing strongly motif-enriched MyoG binding and a shift in Pax7 binding from CpG-rich promoters in progenitors toward Pax7 motif-enriched sites during differentiation. Together, these results show that TF-CUT&Tag captures distinct modes of TF– chromatin interaction, including motif-directed binding sites and more salt-sensitive sites that may depend on lower-affinity DNA recognition or protein–protein interactions and are difficult to predict from DNA sequence alone.

## Introduction

Sequence-specific transcription factors (TFs) coordinate developmental programs and environmental responses by controlling when and where genes are transcribed. The human genome encodes more than 1,600 TFs^1^, but any given cell type expresses only a subset. The expressed TFs then act in a combinatorial manner to select gene promoters, distal enhancers, and other regulatory elements for activation, thereby generating a remarkable degree of regulatory diversity. Predictive models that infer regulatory elements and developmental gene expression from DNA sequence must, at some level, capture how TF combinations select specific genes for transcription, and explicitly defining these regulatory relationships will be necessary to predict and interpret how genetic or environmental perturbations alter genome function in development and disease.

Predicting TF binding from DNA sequence alone remains challenging. Most TFs recognize short, degenerate DNA sequences that occur frequently throughout the genome, but only a fraction of potential recognition sites are occupied in any given cell type^2,3^. Whether a compatible sequence is bound depends on its chromatin and regulatory context, including accessibility, local TF concentration, DNA-binding affinity, and interactions with other TFs and cofactors^3-5^. Consequently, the same TF can bind substantially different loci across cell types, and transcription can depend on the level or frequency of TF binding rather than the simple presence or absence of TF binding^2,6^. Increasing evidence further suggests that TFs can engage chromatin through multiple modes, including direct recognition of high- and low-affinity DNA sequences and cooperative or protein-mediated interactions that can alter both chromatin binding and specificity^7-9^. Thus, DNA sequence provides the foundation for TF recognition, but genome-wide TF binding cannot yet be reliably inferred from sequence alone and must be measured in the relevant cellular context.

Numerous methods have been developed to map TF binding across the genome. Chromatin immunoprecipitation followed by sequencing (ChIP-seq) remains the most widely used method for mapping TF binding. However, ChIP-seq generally requires substantial input material, and the crosslinking and immunoprecipitation steps can introduce artifacts or mask epitopes that limit recovery of some TFs^10,11^. Enzyme-tethering methods provide an alternative strategy by using antibodies to localize DNA-cleaving, modifying, or transposing enzymes to chromatin targets within intact cells or nuclei, thereby mapping protein-DNA interactions *in situ* without chromatin solubilization or immunoprecipitation^12^. For example, antibody-directed MNase cleavage in permeabilized cells substantially reduces CUT&RUN background relative to ChIP-seq, and enables sensitive profiling from small samples and single cells^13,14^. More recently, D&D-seq introduced an antibody-tethered DNA deaminase to label regions of TF binding through localized mutagenesis, allowing TF binding to be coupled with single-cell ATAC-seq and other genomic readouts that preserve cellular context in heterogeneous samples^15^. However, both CUT&RUN and D&D-seq involve tradeoffs between single-cell resolution, genome-wide coverage, and the ability to measure TF binding together with other regulatory features. Although these methods can be applied in complementary settings, their distinct chemistries and readouts introduce different biases in the binding events that are recovered, making it challenging to directly compare and integrate TF binding maps from different assays.

The CUT&Tag method is an attractive, unifying platform because the simple workflow is readily scalable to high-throughput and single-cell applications while also supporting multifactorial measurements. By directing Tn5 transposase to antibody-bound targets, CUT&Tag labels protein-associated DNA directly *in situ* while simultaneously generating sequencing-ready libraries with exceptionally high signal-to-noise^16,17^ (**Fig. 1A**). This workflow has been adapted to automated platforms such as AutoCUT&Tag^18^ and single-cell assays^19,20^. Because Tn5 can be loaded with distinct barcoded adapters, CUT&Tag also supports multifactorial approaches that could, in principle, measure multiple TFs together with histone modifications that provide cellular and chromatin-state context within heterogeneous samples^21-25^. However, realizing this potential has proven difficult because Tn5 has an intrinsic affinity for DNA and requires high-salt washes to suppress non-specific tagmentation at accessible chromatin. These same conditions can destabilize weaker or more transient TF-chromatin interactions, thus limiting sensitivity and potentially distorting the classes of TF binding events recovered in sequencing libraries (**Fig. 1A**).

**Figure 1.**
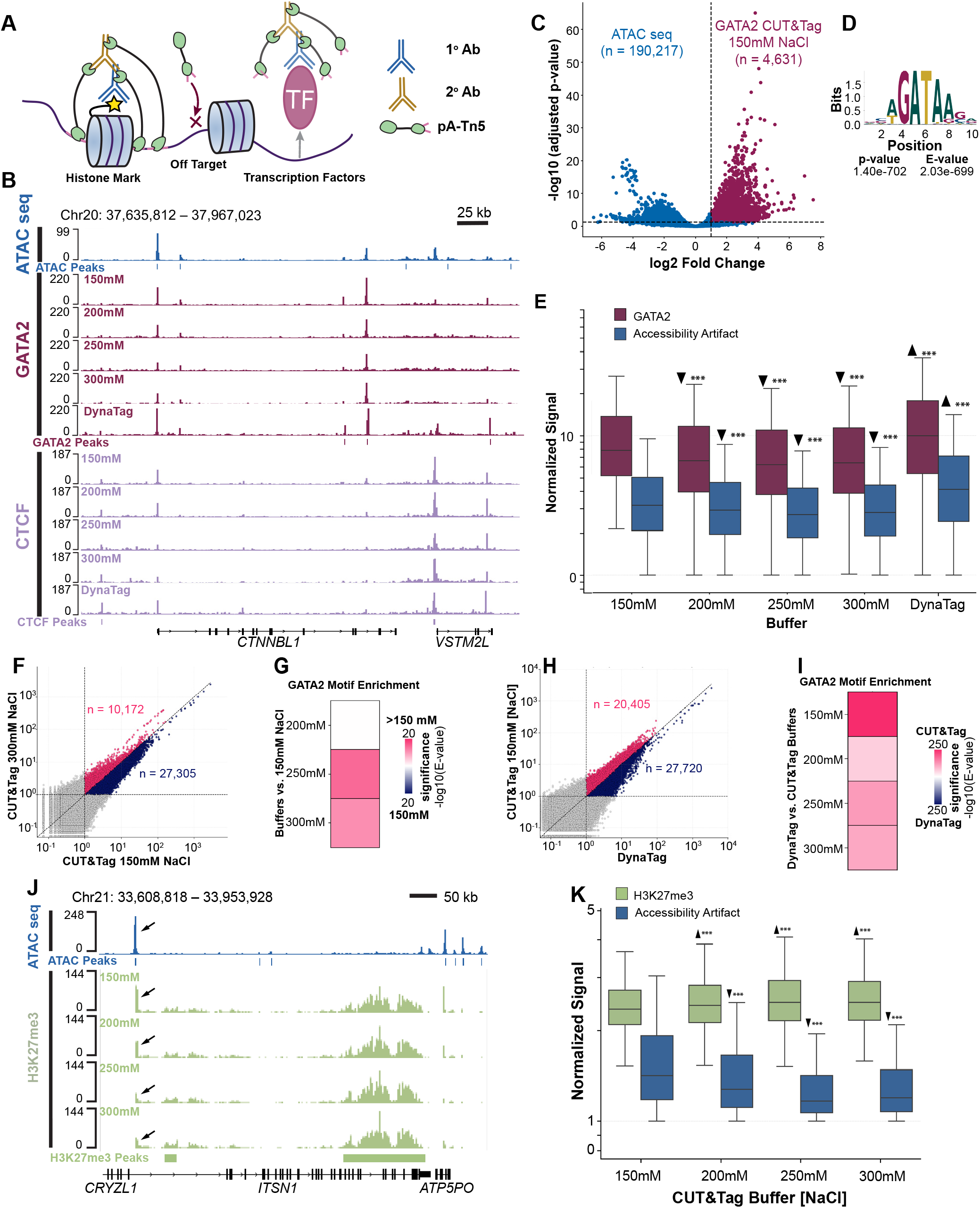
Transcription factor signal and accessibility-associated tagmentation are coupled across ionic conditions. **(A)** CUT&Tag is well suited for profiling histone marks, but the utility for profiling TFs is limited by the high salt washes needed to prevent non-specific Tn5 to off-target sites. **(B)** Genome browser tracks showing ATAC-seq and CUT&Tag profiles of GATA2 and CTCF across the CTNN-BL1 locus at the indicated NaCl concentrations and under DynaTag conditions. **(C)** Volcano plot of DESeq2 results comparing GATA2 CUT&Tag with ATAC-seq. Peaks with log2 fold change > 1 and adjusted P < 0.05 were classified as GATA2-associated peaks (maroon); remaining peaks formed an operational accessibility-associated reference set (blue). **(D)** GATA2 sequence logo and enrichment statistics from Analysis of Motif Enrichment (AME) comparing the GATA2-associated and accessibility-associated peak sets in (C). **(E)** Boxplots showing normalized CUT&Tag signal over GATA2-associated and accessibility-associated sites at the indicated ionic conditions. Boxes show the median and interquartile range (25th–75th percentile), whiskers extend to the most extreme values within 1.5 × the interquartile range, and outliers are not shown. Statistical comparisons are relative to 150 mM NaCl; upward and downward arrowheads indicate the direction of significant changes. **(F**,**H)** Scatterplots comparing normalized signal at merged GATA2 peaks between 150 and 300 mM NaCl (F) or between DynaTag and 150 mM NaCl (H). Each point represents one peak; peaks with signal ≥1 on both axes are colored according to which condition shows higher signal (pink, condition on the y-axis; navy, condition on the x-axis). **(G**,**I)** AME heatmaps showing differential enrichment of the GATA2 motif among the condition-favored peak sets, represented as signed −log10(E-value). In (G), pink indicates enrichment at the indicated NaCl concentration relative to 150 mM NaCl, and blue indicates enrichment at 150 mM NaCl. In (I), pink indicates enrichment under the indicated NaCl condition, and blue indicates enrichment under DynaTag. **(J)** Genome browser tracks showing ATAC-seq and H3K27me3 CUT&Tag across the ITSN1 locus under the indicated NaCl conditions. **(K)** Box-plots of normalized signal over H3K27me3-associated and accessibility-associated peaks across the NaCl gradient. Box-and-whisker elements and statistical symbols are as defined in (E).

Optimizing CUT&Tag can substantially improve the assay’s sensitivity for TF profiling. For example, DynaTag replaces NaCl with physiological concentrations of KCl, which better preserves TF-chromatin interactions^26^. Here, we engineered the Tn5 enzyme itself to suppress untethered transposition at off-target sites of accessible chromatin, allowing us to perform the CUT&Tag assay under lower-salt conditions, which better preserve TF-binding events. We systematically neutralized positively charged residues in the Tn5 DNA-binding channel to reduce intrinsic target-DNA engagement^27^, and identified two variants with TF-dependent effects on the CUT&Tag profiling fidelity. The more strongly attenuated K160A variant improved recovery of GATA2 signal while suppressing untethered tagmentation, but was less effective at profiling CTCF, where the inaccessible local chromatin environment may restrict productive integration. In contrast, K249A selectively improved profiling of CTCF at sites surrounded by tightly spaced nucleosomes, suggesting this variant retains sufficient activity to integrate within narrow windows of accessible DNA. Thus, comparing K160A and K249A provides a new means to assess the physical properties of chromatin surrounding TF-binding sites *in vivo*. Applying TF-CUT&Tag across distinct TFs and cellular states revealed motif-directed and salt-sensitive modes of TF binding, including changes in binding mode during cellular differentiation. These findings demonstrate that engineering Tn5 can suppress off-target tagmentation while retaining TF-chromatin interactions, which will now enable the field to map context-dependent TF binding.

## Results

### Transcription factor signal and off-target tagmentation are coupled across ionic conditions

The ionic stringency that preserves the specificity of histone-mark CUT&Tag may compromise the recovery of salt-sensitive TF–chromatin interactions (**Fig. 1A**). To determine how ionic conditions affect the balance between TF-associated signal and off-target tagmentation, we performed GATA2 and CTCF CUT&Tag with 150, 200, 250, or 300 mM NaCl, and with the DynaTag buffer (110 mM KCl, 10 mM NaCl, and 1 mM MgCl_2_; **Fig. 1B**). Because genuine TF binding sites frequently overlap accessible regulatory elements, we curated a K562 ATAC-seq profile from three publicly available datasets^28-30^, and used this as a reference to distinguish TF-enriched sites from off-target signal. We combined ATAC-seq MACS3 peaks with GATA2 or CTCF CUT&Tag peaks identified with parental pA-Tn5 under 150 mM NaCl conditions, then used DESeq2 to compare the CUT&Tag and ATAC-seq signal. Sites meeting the CUT&Tag-enrichment threshold (log2 fold change > 1 and adjusted *p* < 0.05) were classified as TF-associated peaks, and the remaining sites were defined as the off-target reference set (**Fig. 1C**; **Supplementary Fig. 1A**). This operational reference includes all sites that did not meet the TF-enrichment threshold, rather than only sites significantly enriched in ATAC-seq. Differential motif enrichment analysis supported the distinction between these classes, identifying the cognate GATA2 and CTCF motifs as the most significantly enriched motifs in their respective TF-associated peak sets relative to the off-target reference sets (**Fig. 1D**; **Supplementary Fig. 1B**).

Comparing signals to those from the 150 mM NaCl reference condition revealed that increasing NaCl concentration reduced GATA2-associated and off-target signal (**Fig. 1E**). CTCF showed a similar overall relationship, although both signals initially increased at 200 mM relative to 150 mM before declining at higher NaCl concentrations (**Supplementary Fig. 1C**). Compared with the 150 mM NaCl CUT&Tag conditions, the DynaTag workflow yielded greater normalized GATA2 and CTCF signal but also substantially greater off-target signal (**Fig. 1E**; **Supplementary Fig. 1C**). Thus, conditions yielding greater GATA2-associated signal also yielded greater off-target signal, whereas increasing NaCl concentrations reduced both.

Differential motif enrichment provided an orthogonal measure of this tradeoff. Using a combined peak set of 200bp windows centered on the strongest summits across all CUT&Tag and DynaTag conditions for each TF, we compared those sites with greater normalized signal at 150 mM NaCl against those sites with greater normalized signal at each elevated NaCl concentration (**Fig. 1F**; **Supplementary Fig. 1D–H**). These condition-favored peak sets were defined by the ratio of normalized signal between conditions (greater than or less than 1), rather than by a statistical significance threshold. For both GATA2 and CTCF, the 250 mM-versus-150 mM comparison provided the strongest evidence for motif enrichment in the higher-salt-favored peak set (**Fig. 1G**; **Supplementary Fig. 1I**). These results indicate that sites detected under low salt conditions are comparatively depleted for TF motifs, consistent with reduced off-target tagmentation at elevated ionic strengths.

We next compared sites with greater normalized signal under each NaCl-based CUT&Tag condition with those sites favored by the DynaTag workflow (**Fig. 1H**). For both GATA2 and CTCF, cognate motifs were preferentially enriched in the NaCl-favored peak sets across nearly all comparisons (**Fig. 1I**; **Supplementary Fig. 1J**). The exception was CTCF at 150 mM NaCl, for which there was no significant differential enrichment of the CTCF motif. The lower TF motif enrichment under DynaTag conditions coincided with increased off-target signal. For example, the GATA2 comparison identified 20,405 peaks with greater normalized signal at 150 mM NaCl and 27,720 peaks with greater signal under DynaTag conditions, yet the GATA2 motif was significantly enriched in the smaller 150 mM-favored set (**Fig. 1H,I**). This comparison provided the strongest evidence for GATA2 motif enrichment among the NaCl-versus-DynaTag comparisons (**Fig. 1I**). Together with the signal comparisons, these results indicate that the greater overall TF signal recovered under DynaTag conditions coincides with increased off-target tagmentation. To determine if higher NaCl concentrations also reduced on-target signal for a histone modification, we profiled H3K27me3 across the same NaCl gradient. Examining a representative genomic region revealed prominent signal at accessible chromatin under 150 mM NaCl, a signal that progressively diminished as NaCl concentration increased (**Fig. 1J**). In contrast, normalized signal within H3K27me3-marked domains was retained at higher NaCl concentrations (**Fig. 1J**). Genome-wide quantification using H3K27me3 peaks called at 300 mM NaCl and the non-overlapping portion of the curated K562 ATAC-seq peak set confirmed that increasing NaCl concentrations reduced off-target signal while retaining normalized H3K27me3 signal (**Fig. 1K**). Thus, increasing ionic strength suppressed off-target tagmentation without decreasing normalized H3K27me3 on-target signal. Taken together, these results show that increasing ionic strength suppresses off-target tagmentation at the cost of TF-associated signal, consistent with the loss of salt-sensitive TF–chromatin interactions

### Reducing positive charge in the Tn5 DNA-binding channel suppresses off-target tagmentation

The coupled changes in TF-associated and off-target signal motivated us to alter Tn5 itself rather than relying solely on buffer optimization. We therefore asked if we could reduce the intrinsic Tn5 DNA binding affinity while preserving antibody-tethered tagmentation. Because the Tn5 DNA-binding channel contains positively charged residues that electrostatically interact with DNA, we reasoned that neutralizing these contacts would preferentially reduce untethered DNA engagement at accessible chromatin, yet retain productive transposition when Tn5 is localized to its target via antibody tethering (**Fig. 2A**).

**Figure 2.**
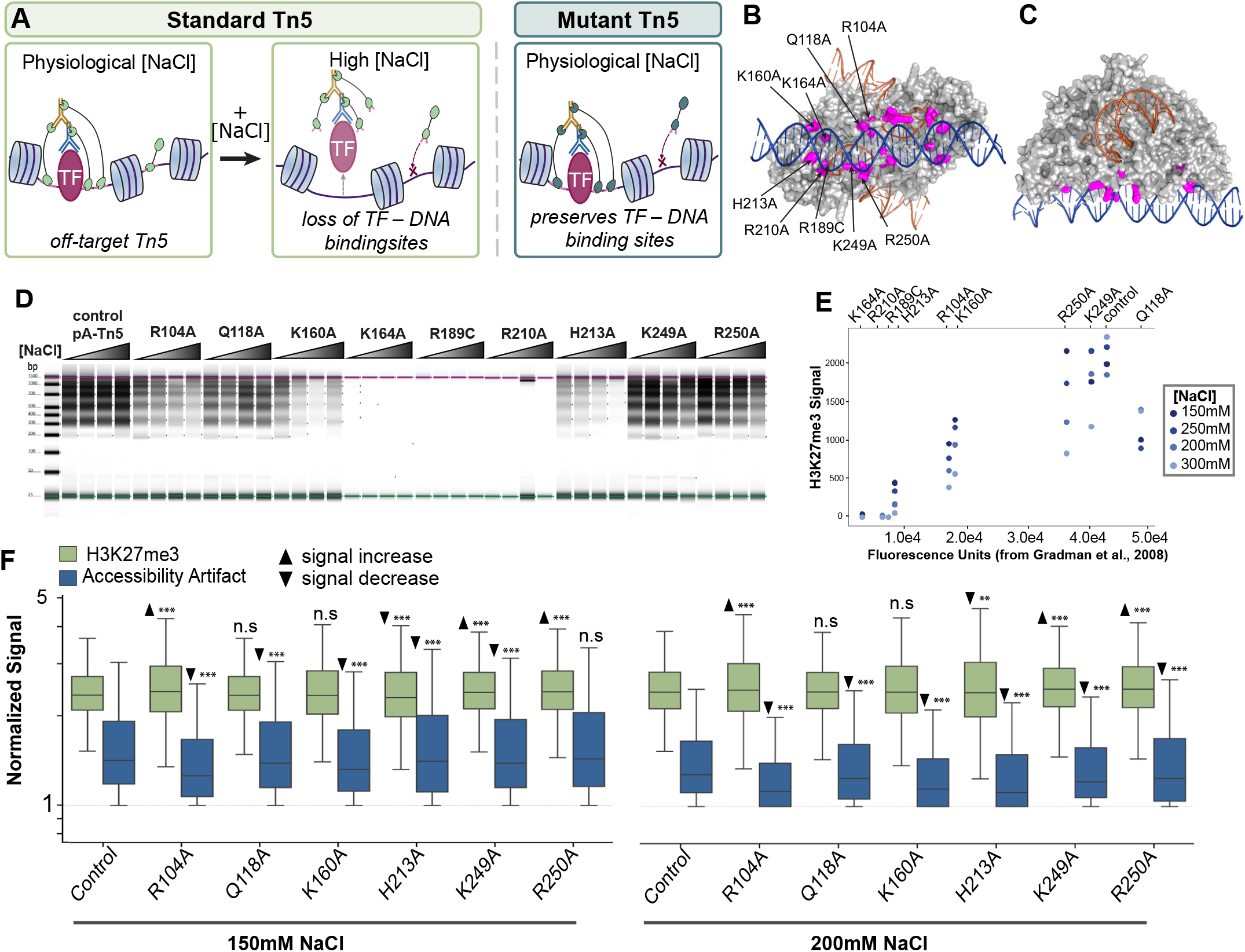
Reducing positive charge in the Tn5 DNA-binding channel suppresses off-target tagmentation. **(A)** Schematic illustrating the rationale for modifying Tn5 to suppress untethered insertion into accessible chromatin while preserving antibody-directed tagmentation at TF-bound sites under lower-stringency ionic conditions. **(B**,**C)** Structural models of Tn5 associated with DNA, viewed from two orientations, showing the positions of residues selected for mutagenesis (magenta). **(D)** Bioanalyzer virtual-gel profiles of H3K27me3 CUT&Tag libraries generated with parental pA-Tn5 or the indicated variants at 150, 200, 250, and 300 mM NaCl; ionic concentration increases from left to right within each group. **(E)** Comparison of H3K27me3 CUT&Tag signal with previously reported Tn5 target-capture/strand-transfer activity (fluorescence units; Gradman et al., 2008). Points represent individual variant and buffer combinations. **(F)** Boxplots of normalized H3K27me3-associated (green) and accessibility-associated (blue) signal for parental pA-Tn5 and the indicated variants at 150 and 200 mM NaCl. Box-and-whisker elements are as defined in Fig. 1E. Statistical comparisons are between each variant and parental pA-Tn5 within the same salt condition, with upward and downward arrowheads indicating the direction of significant changes.

Guided by previous mutational analyses of Tn5 target capture and strand-transfer activity^27^, we generated a panel of pA-Tn5 variants carrying the substitutions R104A, Q118A, K160A, K164A, R189C, R210A, H213A, K249A, or R250A (**Fig. 2B**). Each of these residues contributes to DNA binding and catalytic activity, and converting the positively charged residues to alanine allowed us to test if reduced electrostatic engagement with DNA could suppress off-target tagmentation without eliminating the Tn5 catalytic activity required for CUT&Tag. Importantly, the mutations did not substantially alter the Tn5 loading efficiency with annealed adapter oligonucleotides, indicating that differences in CUT&Tag activity among the variants were not attributable to differences in transposome assembly (**Supplementary Fig. 2A–C**).

To evaluate retained on-target activity and off-target tagmentation for each variant, we first profiled H3K27me3, for which signal over H3K27me3-associated peaks could be assessed separately from the off-target reference peaks. The K164A, R189C, and R210A variants produced little or no detectable H3K27me3 signal across the tested conditions, indicating that these substitutions substantially impaired productive CUT&Tag (**Fig. 2D**). R104A, K160A, and H213A variants retained intermediate activity but progressively lost H3K27me3 signal at increasing NaCl concentrations (**Fig. 2D**), indicating these substitutions sensitize Tn5 activity to salt concentration. In contrast, Q118A, K249A, and R250A variants produced H3K27me3 signal approaching that of parental pA-Tn5 (**Fig. 2D**), with the variation in H3K27me3 signal broadly reflecting previously reported differences in the activity of these same mutants in a biochemical target-capture/strand-transfer assay^27^ (**Fig. 2E**). Thus, mutating positively charged amino acids in the Tn5 DNA binding channel had varied effects on CUT&Tag activity, with a subset of variants exhibiting pronounced sensitivity to ionic conditions.

Having established which variants retained productive CUT&Tag activity, we next asked if any of these variants also reduced off-target tagmentation. Multiple variants that retained H3K27me3 signal showed reduced signal at off-target sites under lower-salt conditions (**Fig. 2F, Suppl. Fig 2D**). This effect was most apparent at 150 and 200 mM NaCl, where the parental enzyme exhibited substantial off-target tagmentation. Thus, some mutations in the Tn5 DNA binding channel can suppress off-target tagmentation while preserving productive antibody-directed activity under the same buffer conditions.

### Engineered Tn5 variants show TF-specific effects on CUT&Tag fidelity

To determine if reduced off-target tagmentation improved CUT&Tag performance for TFs, we profiled GATA2 and CTCF using the pA-Tn5 variants that retained enzymatic activity. At 150 mM NaCl, K160A was the only variant that significantly increased GATA2-associated signal while simultaneously reducing off-target signal relative to parental pA-Tn5 (GATA2-associated sites, p = 2.38 × 10^-30^; off-target sites, p = 5.11 × 10^-4^; **Fig. 3A**). Other variants that increased GATA2-associated sig GATA2-associated signal, although off-target signal also increased (p = 1.96 × 10^-71^ and p = 2.86 × 10^-6^, respectively). At 200 mM NaCl, H213A also increased GATA2-associated signal and reduced off-target signal relative to parental pA-Tn5 (**Fig. 3A**). Thus, K160A is the only variant that improved the balance between GATA2-associated and off-target signal at 150 mM NaCl, while the H213A variant achieves a similar separation at 200 mM NaCl.

**Figure 3.**
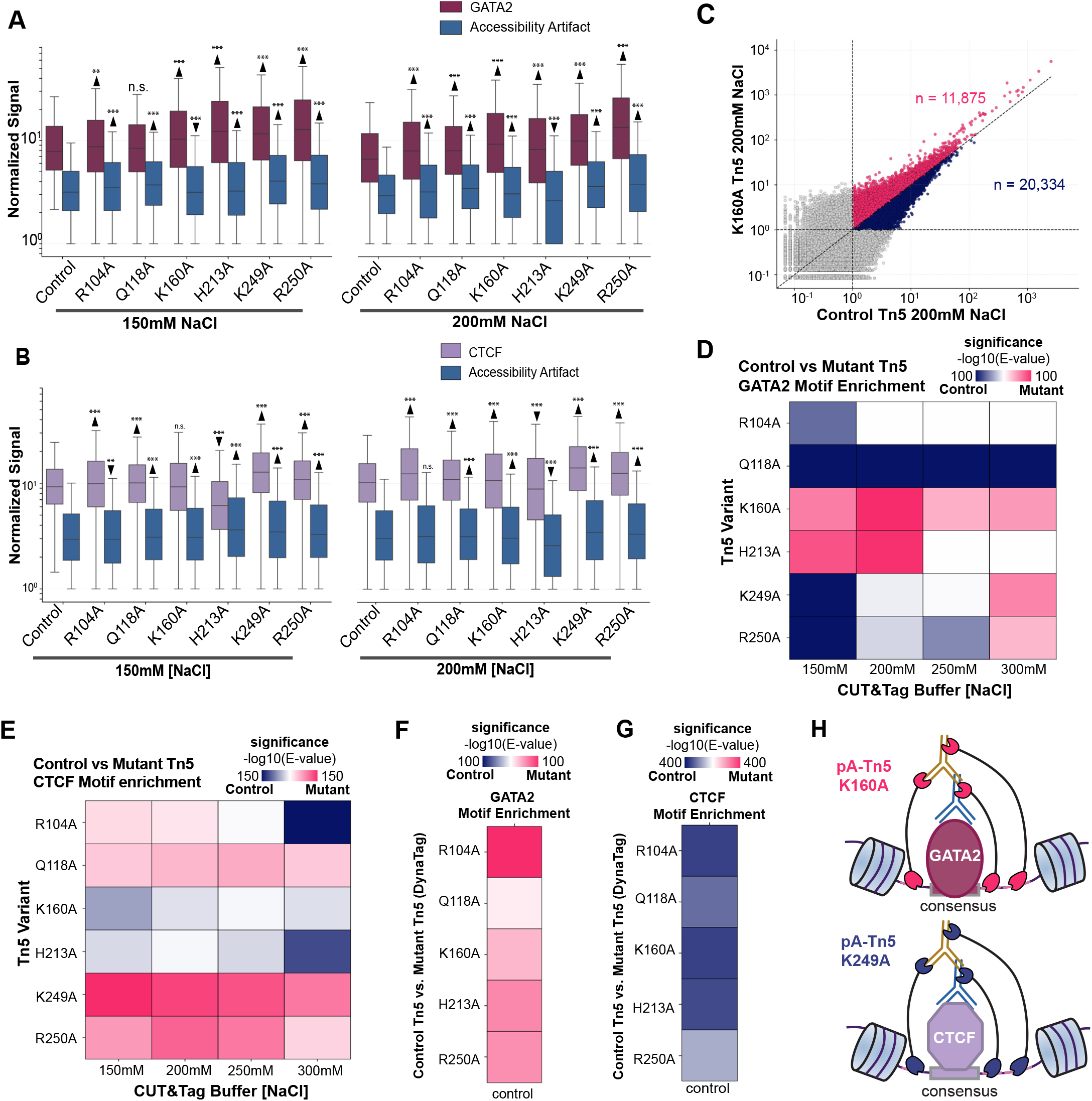
Engineered Tn5 variants show TF-specific effects on CUT&Tag fidelity. **(A,B)** Boxplots of normalized GATA2-associated and accessibility-associated signal (A) or CTCF-associated and accessibility-associated signal (B), comparing parental pA-Tn5 with the indicated variants at 150 and 200 mM NaCl. TF-associated and accessibility-associated sites were defined as in Fig. 1C and Supplementary Fig. 1A, respectively. Box-and-whisker elements are as defined in Fig. 1E; statistical comparisons are between each variant and parental pA-Tn5 at matched NaCl concentrations, with arrowheads indicating the direction of significant changes. **(C)** Scatterplot comparing normalized GATA2 CUT&Tag signal for K160A and parental pA-Tn5 at 200 mM NaCl across the merged GATA2 peak set. Each point represents a peak; among peaks with signal ≥1 on both axes, pink indicates K160A-favored sites and navy indicates parental-enzyme-favored sites. **(D**,**E)** Heatmaps showing differential enrichment of cognate GATA2 (D) or CTCF (E) motifs by AME between variant-favored and parental-enzyme-favored peak sets under matched NaCl conditions, expressed as signed −log10(E-value). Pink indicates enrichment among variant-favored peaks; blue indicates enrichment among parental-enzyme-favored peaks. **(F**,**G)** Corresponding AME comparisons of each indicated variant with parental pA-Tn5 under DynaTag conditions for GATA2 (F) and CTCF (G). Color direction is as in (D,E). **(H)** Schematic summarizing the contrasting TF-dependent performance of engineered pA-Tn5: K160A favors recovery of GATA2-associated sites, whereas K249A performs well at CTCF-associated sites. The schematic represents a proposed interpretation rather than a direct measurement of Tn5–DNA affinity.

We next performed a similar comparison over the CTCF sites. In contrast to GATA2, K160A and H213A did not selectively improve CTCF-associated signal relative to the parental Tn5 (**Fig. 3B**). Instead, K249A and R250A increased signal at CTCF-associated peaks, but these increases were accompanied by greater signal at off-target sites. These different responses suggested that the degree of Tn5 attenuation favorable for one TF might not improve recovery of another.

To determine if these signal changes were associated with preferential recovery of motif-containing sites, we compared normalized signal across a merged GATA2 peak set assembled from all conditions (The comparison at 200mM NaCl is shown as an example; **Fig. 3C**). We then performed differential motif enrichment analysis on peak sets favoring each variant or parental pA-Tn5 under identical ionic conditions (**Fig. 3D**). At 150 and 200 mM NaCl, K160A-favored peaks showed the strongest evidence for GATA2 motif enrichment relative to parental-enzyme-favored peaks, followed by the H213A variant (**Fig. 3D**). Together with the signal measurements, these results support a genuine improvement in GATA2 profiling at 150 mM and 200 mM NaCl concentrations.

Across variants, the CTCF motif analysis revealed a near mirror image of the GATA2 motif analysis (**Fig. 3E**). In contrast to GATA2 profiling, the K160A and H213A variants showed a reduction in CTCF motif enrichment relative to parental pA-Tn5. Conversely, peaks favored by K249A and R250A were consistently enriched for the CTCF motif (**Fig. 3E**), even though these variants were less effective for GATA2. This reversal in performance suggests the preferred variant for TF-CUT&Tag may vary depending on the target.

To determine whether technical artifacts could explain the opposing GATA2 and CTCF results, we first examined the fraction of reads in peaks (FRiP). Most variants increased FRiP relative to parental pA-Tn5 across salt conditions, particularly K249A and R250A for both factors (**Supplementary Fig. 3A,B**). Because differences in FRiP can influence peak comparisons after whole-library normalization (**Supplementary Fig. 3C,D**), we repeated the motif analysis after normalizing to fragments within the consensus peak set. The TF-dependent relationships persisted, with K160A- and H213A-favored peaks enriched for GATA2 motifs, whereas K249A- and R250A-favored peaks were enriched for CTCF motifs (**Supplementary Fig. 3E,F**). Tn5 insertion-site sequence preferences were also broadly similar across parental and mutant enzymes, with only a modest change for K160A at GATA2 sites (**Supplementary Fig. 3G**). Thus, neither normalization effects nor altered Tn5 insertion preferences readily explained the opposing motif-enrichment patterns.

Potassium can stabilize nucleosomal DNA wrapping relative to sodium *in vitro*^*31*^, and to test whether physical differences in the local chromatin environment at GATA2 and CTCF sites might underlie the variant-to-TF relationships we observed, we compared the variants to parental Tn5 under potassium-rich DynaTag conditions. Across GATA2 profiles, the peaks favored by the Tn5 variants were enriched for GATA2 motifs relative to peaks favored by parental pA-Tn5 (**Fig. 3F**). In contrast, CTCF motifs were preferentially enriched among peaks favored by parental pA-Tn5 rather than any of the engineered variants (**Fig. 3G**). These opposing responses are consistent with differences in the physical organization of chromatin surrounding GATA2 and CTCF sites, where stabilizing the nucleosomes at more restricted CTCF sites could limit productive integration by the attenuated Tn5 variants. Notably, direct comparisons of each enzyme between NaCl-based CUT&Tag and DynaTag generally showed greater CTCF and GATA2 motif enrichment in the peaks favored under NaCl conditions (**Supplementary Fig. 3H,I**).

Together, these experiments identify K160A as the most effective variant for preferential recovery of GATA2 motif-containing sites at relatively low NaCl concentrations, whereas K249A is better suited to profiling CTCF (**Fig. 3H**).

### Fragment architecture and accessibility distinguish the TF-binding sites preferred by K160A and K249A

The differential responses of the Tn5 variants under DynaTag suggests that the chromatin environment surrounding the targeted TF influences the amount of residual Tn5 DNA-engagement activity required for productive tagmentation. To investigate the physical differences that may underlie this observation, we directly compared the K249A and K160A variants at 200 mM NaCl. For both GATA2 and CTCF, we quantified signal across TF-specific sets of 200-bp consensus peaks and plotted the log2(K160A/K249A) signal ratio against the strongest cognate-motif score within each peak (**Fig. 4A,B**). We then defined two opposing peak sets for each factor, pairing the direction of mutant preference with higher or lower motif scores. For GATA2, these were high-motif, K160A-favored peaks (n = 8,311) and lower-motif, K249A-favored peaks (n = 8,312). For CTCF, the relationship was reversed, and high-motif K249A-favored peaks (n = 14,485), and lower-motif K160A-favored peaks (n = 14,485) were selected. Motif-score thresholds were chosen separately for each TF to balance group sizes (**Supplementary Fig. 4A,B**). These intentionally contrasting peak populations were used as test cases to investigate what properties of TF-associated chromatin might account for the different variant preferences.

**Figure 4.**
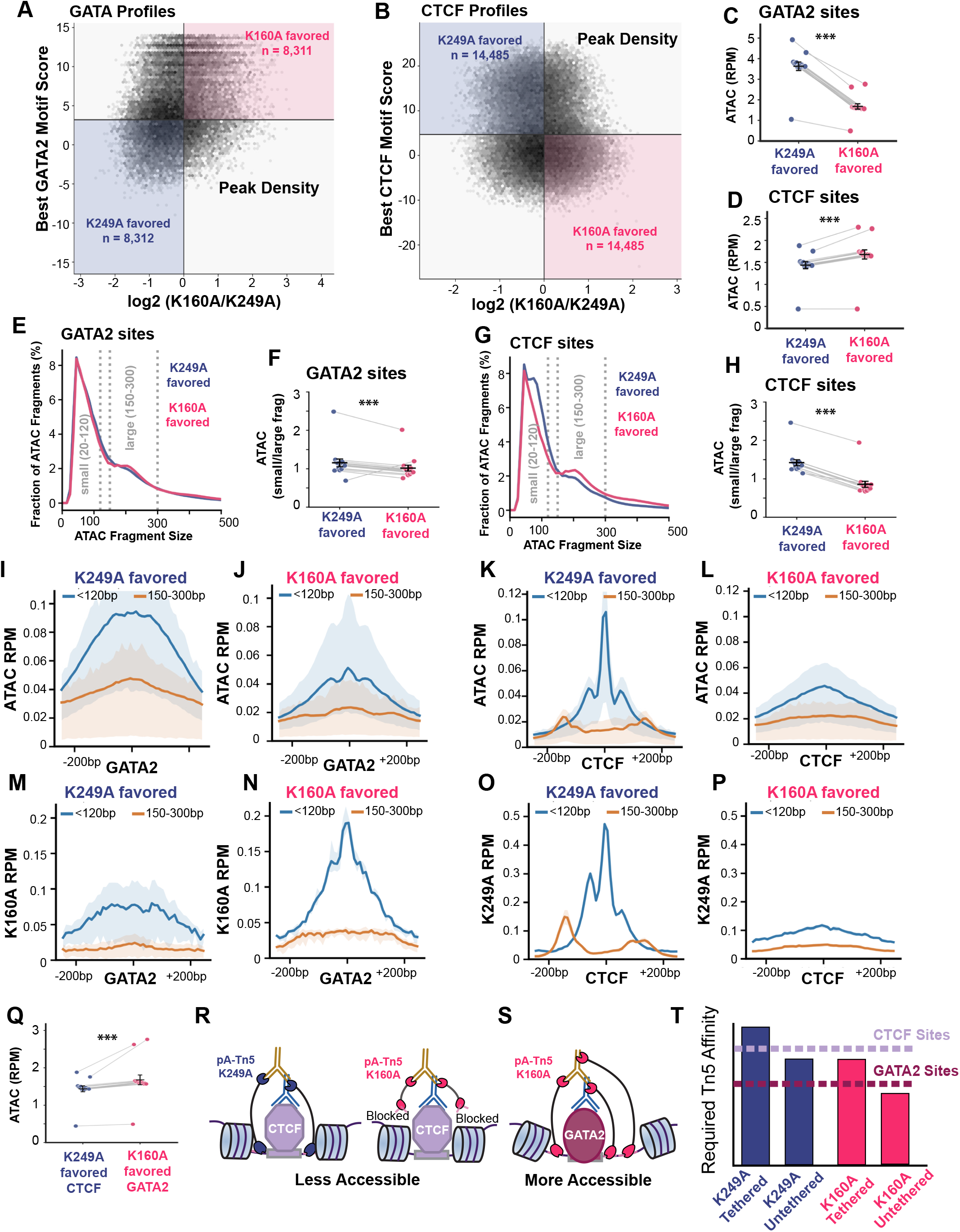
Fragment architecture and accessibility distinguish the TF-binding sites preferred by K160A and K249A. **(A,B)** Density plots comparing the best-scoring cognate motif within each 200-bp consensus peak with the log2 ratio of normalized K160A/ K249A CUT&Tag signal at 200 mM NaCl, shown for GATA2 (A) and CTCF (B). Selected opposing quadrants identify K160A-preferred, high-motif GATA2 sites and K249A-preferred, low-motif GATA2 sites (A), or K249A-preferred, high-motif CTCF sites and K160A-preferred, low-motif CTCF sites (B). Motif-score thresholds were selected to yield balanced preferred populations (Supplementary Fig. 4A,B); other sites are shown in gray. **(C**,**D)** Paired-library comparisons of mean ATAC-seq fragment RPM per peak at the selected K249A- and K160A-preferred GATA2 (C) and CTCF (D) sites. Each point denotes one ATAC-seq library; lines connect measurements from the same library, and black symbols indicate the mean ± SEM. **(E**,**F)** ATAC-seq fragment-length distributions for the selected GATA2 (E) and CTCF (F) populations. Dashed lines delimit the short (20–119 bp) and nucleosome-sized (150–299 bp) fragment intervals. **(G**,**H)** Paired-library comparisons of ATAC-seq short-to-nucleosome-sized fragment ratios across the GATA2 (G) and CTCF (H) peak populations. **(I–L)** Motif-centered mean ATAC-seq fragment-midpoint profiles for the K249A- and K160A-preferred GATA2 populations (I,J) and CTCF populations (K,L). Blue indicates short fragments (20–119 bp) and orange indicates nucleosome-sized fragments (150–299 bp); shaded envelopes show the range across individual libraries. (M,N; left pair) Corresponding motif-centered K160A CUT&Tag profiles at the GATA2 populations. (O,P; right pair) Corresponding motif-centered K249A CUT&Tag profiles at the CTCF populations. **(Q)** Paired ATAC-seq comparison between K249A-preferred CTCF sites and K160A-preferred GATA2 sites, using the same ATAC libraries for both peak populations. **(R**,**S)** Proposed model in which restricted DNA accessibility around CTCF limits productive integration by the more strongly attenuated K160A variant, whereas antibody recruitment permits K160A integration near GATA2-associated sites with greater local DNA availability. **(T)** Conceptual model of the effective DNA-engagement activity required for productive tagmentation at the two chromatin environments, comparing tethered and untethered K249A and K160A. Bars and threshold lines are illustrative, not biochemical affinity measurements.

To examine physical differences at GATA2 and CTCF binding sites, We first used independently generated ATAC-seq libraries to compare the accessibility of these selected populations. For GATA2, the higher-motif sites preferentially recovered by K160A had significantly lower average ATAC signal than the lower-motif, K249A-preferred sites (**Fig. 4C**). For CTCF, the higher-motif sites preferred by K249A were likewise less ATAC-accessible than the K160A-preferred sites (**Fig. 4D**). Thus, in both cases the variant that better recovered cognate-motif-associated peaks favored a peak set with lower conventional ATAC accessibility. This is consistent with antibody tethering enabling productive tagmentation at factor-associated regions across the genome that are relatively inaccessible to ATAC-seq.

We next examined absolute CUT&Tag recovery across these same peak sets. Parental pA-Tn5 showed little separation between the two GATA2 populations, whereas K160A tended to recover more signal from its preferred, high-motif GATA2 sites (**Supplementary Fig. 4C,D**). At CTCF sites, K249A showed greater mean signal in its preferred high-motif population, with different degrees of separation in the parental and K160A profiles (**Supplementary Fig. 4E,F**). These results indicate that the variant-specific preferences are reflected in differences in the mean CUT&Tag signal recovered at these sites, rather than arising solely from the ratio between K160A and K249A used to select them.

We then asked whether the Tn5-variant-preferred binding sites differed in fragment organization. ATAC-seq fragment-size distributions showed relatively modest differences between the GATA2 populations (**Fig. 4E,F**), but a much more pronounced excess of short (20–120 bp) relative to nucleosome-sized (150–300 bp) fragments at K249A-preferred CTCF sites (**Fig. 4G,H**). The enrichment of short fragments at these CTCF sites was also observed in parental and mutant CUT&Tag libraries, not only in the K249A profiles (**Supplementary Fig. 4G-I**). This suggested that the short-fragment bias reflected a property of these selected chromatin regions rather than an intrinsic preference of K249A for producing a particular fragment size.

To examine the spatial organization of proteins over these sites, we next centered the short- and nucleosome-sized fragments on the best-scoring GATA2 or CTCF motif in each peak. ATAC-seq profiles showed an accumulation of short-fragment midpoints around the cognate motif for both factors, with differences in their spatial distribution between the K249A and K160A selected peak sets (**Fig. 4I–L**). At K160A-preferred GATA2 sites, short fragments formed three local peaks surrounding the central GATA2 motif. In contrast, nucleosome-sized fragments showed a more diffuse distribution, with a central enrichment consistent with a positioned nucleosome and two weaker regions of enrichment flanking the region enriched for short fragments. The distribution of small and large fragments was even more accentuated at CTCF sites. Here, K249A-favored peaks exhibited a pronounced sharp peak directly over the CTCF motif, and two less pronounced peaks on either side, followed by an accumulation of nucleosomal fragments (**Fig. 4K**), suggesting these CTCF bound sites are flanked by tightly positioned nucleosomes. In comparison, K160A-favored peaks showed a much broader and weaker central enrichment (**Fig. 4L**). Thus, the sites preferred by the two variants differed not only in overall accessibility but in the geometry of protein mediated protections and nucleosome spacing near the predicted TF-binding site.

The motif-centered CUT&Tag profiles reinforced this interpretation. K160A produced a pronounced central short-fragment signal at its preferred high-motif GATA2 sites (**Fig. 4M,N)**, whereas K249A generated a particularly sharp short-fragment peak over the motifs in its favored CTCF peak set (**Fig. 4O,P**). The same general patterns were also detectable when K249A was applied to GATA2 and K160A to CTCF, but were less pronounced in the populations preferred by the opposing variant (**Supplementary Fig. 4K-N**). These data are consistent with K249A retaining a greater capacity for productive integration within narrow, spatially restricted windows of accessible DNA that fall between tightly bound CTCF proteins (represented by the central small fragment peaks), and flanking nucleosomes that directly abut CTCF. K160A can recover such sites, but its greater attenuation may limit integration even when tethered.

A central prediction of this model is that the K249A-preferred CTCF sites should be less accessible than the GATA2 sites efficiently recovered by K160A. Directly comparing the same ATAC-seq libraries across these two selected populations showed that, indeed, the K249A-preferred CTCF sites had significantly lower mean ATAC signal than the K160A-preferred GATA2 sites (**Fig. 4Q**). Together, these findings support a model in which successful TF-CUT&Tag integration depends on the balance between residual Tn5 DNA-engagement activity, antibody-mediated recruitment, and the availability of nearby accessible DNA for insertion (**Fig. 4P,Q**). K249A may retain sufficient activity to engage accessible DNA within the more confined CTCF-associated chromatin environment, whereas K160A may be too strongly attenuated to integrate efficiently at some of these sites, even when tethered (**Fig. 4R**). At the more permissive GATA2-associated sites, however, K160A appears to retain enough activity for recruitment-dependent insertion while reducing untethered accessibility-associated tagmentation (**Fig. 4S**). Thus, engineering Tn5 can improve the balance between TF-directed recovery and accessibility-associated background under relatively low-salt conditions, but the optimal degree of attenuation depends on the local chromatin environment (**Fig. 4T**). We conclude that comparing K249A and K160A TF-CUT&Tag profiles of the same TF provides a new approach to assess the physical chromatin environment at TF binding sites.

### TF-CUT&Tag improves recovery of pluripotency transcription factor binding

Because high fidelity CTCF sites can be obtained by standard CUT&Tag at higher NaCl concentrations^32^, we next asked whether K160A could extend the benefits of low-salt TF profiling across additional, potentially more salt-sensitive transcription factors. We profiled the pluripotency-associated factors Oct4, Sox2, Nanog, and cMyc in mouse embryonic stem cells using TF-CUT&Tag with K160A at 200 mM NaCl, and standard CUT&Tag with parental pA-Tn5 at 300 mM NaCl. Around the *Nanog* locus, TF-CUT&Tag produced stronger signal and retained the overall pattern of TF binding observed with standard CUT&Tag for all TFs (**Fig. 5A**). Genome-wide, TF-CUT&Tag increased the FRiPs for each factor relative to standard CUT&Tag, with matched IgG controls showing lower FRiP values (**Fig. 5B-E**). Peak calling revealed a substantial increase in recovered sites for Oct4 and Nanog, with 2,662 peaks called with standard CUT&Tag for Oct4 versus 4,946 Oct4 peaks with TF-CUT&Tag, and 6,619 Nanog peaks with standard CUT&Tag versus 11,055 Nanog peaks with TF-CUT&Tag. In contrast, the total numbers of Sox2 and cMyc peaks were nearly unchanged between standard CUT&Tag and TF-CUT&Tag (9,096 versus 9,129 for Sox2 and 4,101 versus 4,133 for cMyc, respectively). Thus, TF-CUT&Tag improved overall recovery for all four factors, although the extent to which this translated into additional called peaks differed by TF.

**Figure 5.**
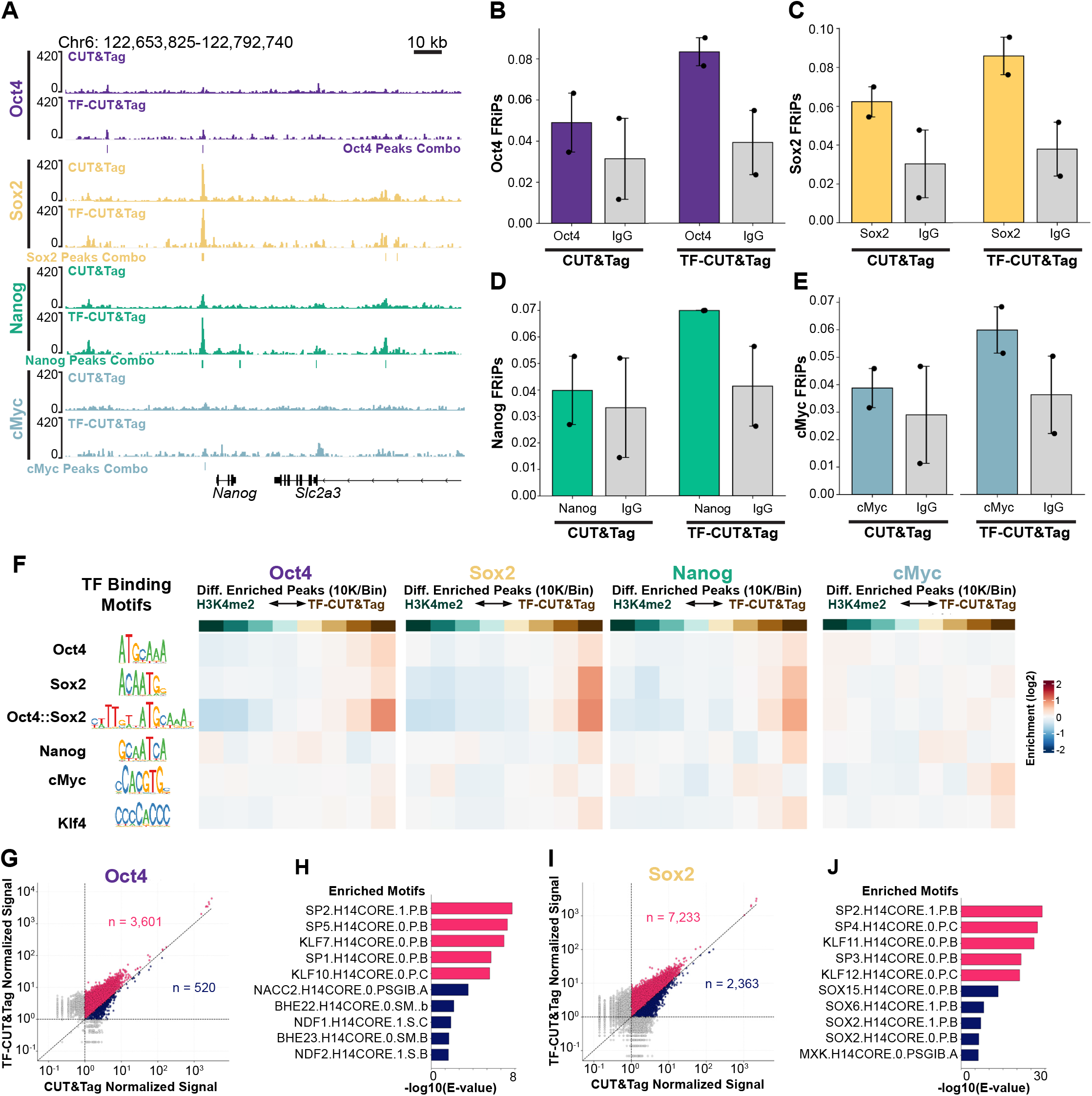
TF-CUT&Tag improves recovery of pluripotency transcription factor occupancy. **(A)** Genome browser tracks showing Oct4, Sox2, Nanog, and cMyc profiles in mouse embryonic stem cells generated by standard CUT&Tag with parental pA-Tn5 at 300 mM NaCl or TF-CUT&Tag with K160A at 200 mM NaCl across the Nanog locus. **(B–E)** Fraction of reads in peaks (FRiP) for Oct4 (B), Sox2 (C), Nanog (D), and cMyc (E) in standard CUT&Tag and TF-CUT&Tag, together with matched IgG controls. Bars summarize replicate measurements; black dots indicate individual libraries. **(F)** Heatmaps showing monaLisa enrichment of the indicated pluripotency-associated motifs across consecutive 10,000-site bins ordered by relative enrichment between H3K4me2 CUT&Tag and TF-CUT&Tag for each TF, from H3K4me2-favored to TF-CUT&Tag-favored sites. **(G**,**I)** Scatterplots comparing normalized signal across merged peak sets between TF-CUT&Tag and standard CUT&Tag for Oct4 (G) and Sox2 (I). Each point represents one peak; among peaks with normalized signal ≥1 on both axes, pink indicates TF-CUT&Tag-favored sites and navy indicates standard-CUT&Tag-favored sites. **(H**,**J)** Bar plots showing differential AME motif enrichment between TF-CUT&Tag-favored and standard-CUT&Tag-favored peaks for Oct4 (H) and Sox2 (J), expressed as −log10(E-value). Pink and blue bars indicate motifs enriched among TF-CUT&Tag- and stan-dard-CUT&Tag-favored sites, respectively.

We next examined the sequence features associated with sites preferentially recovered by TF-CUT&Tag. Because H3K4me2 marks active promoters and enhancers and broadly overlaps accessible regulatory chromatin^33^, we used H3K4me2 CUT&Tag as a reference for regulatory-element-associated signal. We ranked sites by their relative enrichment for each TF versus H3K4me2, and used monaLisa to assess motif enrichment across consecutive 10,000-site bins (**Fig. 5F**). Oct4 and Sox2 profiles showed enrichment for Oct4-, Sox2-, and composite Oct4::Sox2-associated motifs toward the TF-enriched end of the distribution. Nanog-enriched sites likewise showed enrichment for Sox2 and composite Oct4::Sox2 motifs, whereas cMyc showed a weaker and distinct pattern across the motifs examined. These data show that the sequence features enriched by TF-CUT&Tag differed among factors, with particularly strong cognate or cooperative motif enrichment for Oct4, Sox2, and Nanog.

We then compared normalized signal between TF-CUT&Tag and standard CUT&Tag across a merged peak set generated for each factor. TF-CUT&Tag produced higher signal across more sites relative to standard CUT&Tag for all four factors (**Fig. 5G, I; Supplementary Fig. 5I,K**). Thus, TF-CUT&Tag increased signal across thousands of sites for which the total numbers of called peaks were nearly unchanged. However, these signal gains were not uniformly accompanied by greater cognate-motif enrichment in the TF-CUT&Tag-favored peak sets. For example, pairwise motif analysis did not identify the cognate Oct4, Nanog, or cMyc motifs as preferentially enriched in either condition-favored peak set, whereas the Sox2 motif was more enriched among sites favored by standard CUT&Tag (**Fig. 5H,J; Supplementary Fig. 5J,L)**. Similar to CTCF, Sox2 therefore showed stronger cognate-motif enrichment among sites favored by the standard (more stringent) CUT&Tag condition, despite an overall increase in signal with TF-CUT&Tag. Together, these results show that TF-CUT&Tag improves signal recovery across multiple pluripotency-associated TF profiles, but that the relationship between increased recovery and cognate-motif enrichment differs among factors.

### TF-CUT&Tag reveals distinct MyoG and Pax7 binding modes during myogenic differentiation

Having established improved TF specificity and signal recovery with K160A, we next asked if TF-CUT&Tag could detect dynamic TF binding during cellular differentiation. C2C12 myogenic differentiation provides a well-characterized transition in which Pax7 is associated with the progenitor state, whereas MyoG is induced as cells commit to differentiation^34^. Once expressed, MyoG binds and transactivates the terminal myogenic program, including myosin genes and regulatory factors such as Myomixer (Mymx). We profiled both factors using standard CUT&Tag and TF-CUT&Tag in proliferative, undifferentiated myoblasts (Day -1; 24 hours before confluence and switch to differentiation media) and after 24 or 48 hours in differentiation media (Day 1 and Day 2), with matched IgG controls at each time point (**Fig. 6A**). Using the Mymx locus as a well characterized MyoG target gene, TF-CUT&Tag detected little MyoG signal at Day -1, weak signal at Day 1, and prominent enrichment at Day 2, whereas standard CUT&Tag recovered substantially weaker signal across the same time interval (**Fig. 6B**). Genome-wide quantification using a merged Day 2 reference peak set showed that MyoG FRiP was highest at Day 2 and was 3.22-fold greater with TF-CUT&Tag than with standard CUT&Tag, while IgG FRiP remained substantially below the MyoG-directed signal (**Fig. 6C**). We then compared Day 2 MyoG TF-CUT&Tag with H3K4me2 CUT&Tag across a merged peak set (**Fig. 6D**). Ranking sites by their relative enrichment for MyoG revealed progressively stronger enrichment of myogenic E-box motifs toward the MyoG-enriched end of the distribution, including motifs annotated to Myf6, MyoD1, and MyoG (**Fig. 6E**). This pattern is consistent with sequence-directed MyoG binding at myogenic regulatory elements. Comparing the two assays also showed substantially more sites with higher normalized signal in TF-CUT&Tag relative to standard CUT&Tag (**Fig. 6F**). Sites favored by TF-CUT&Tag were preferentially enriched for myogenic E-box motifs annotated to MyoD, Myf5, and related basic helix-loop-helix motifs, whereas sites favored by standard CUT&Tag showed enrichment for a different set of motifs (**Fig. 6G**). Thus, the increased recovery of MyoG signal included preferential recovery of sites bearing myogenic recognition sequences.

**Figure 6.**
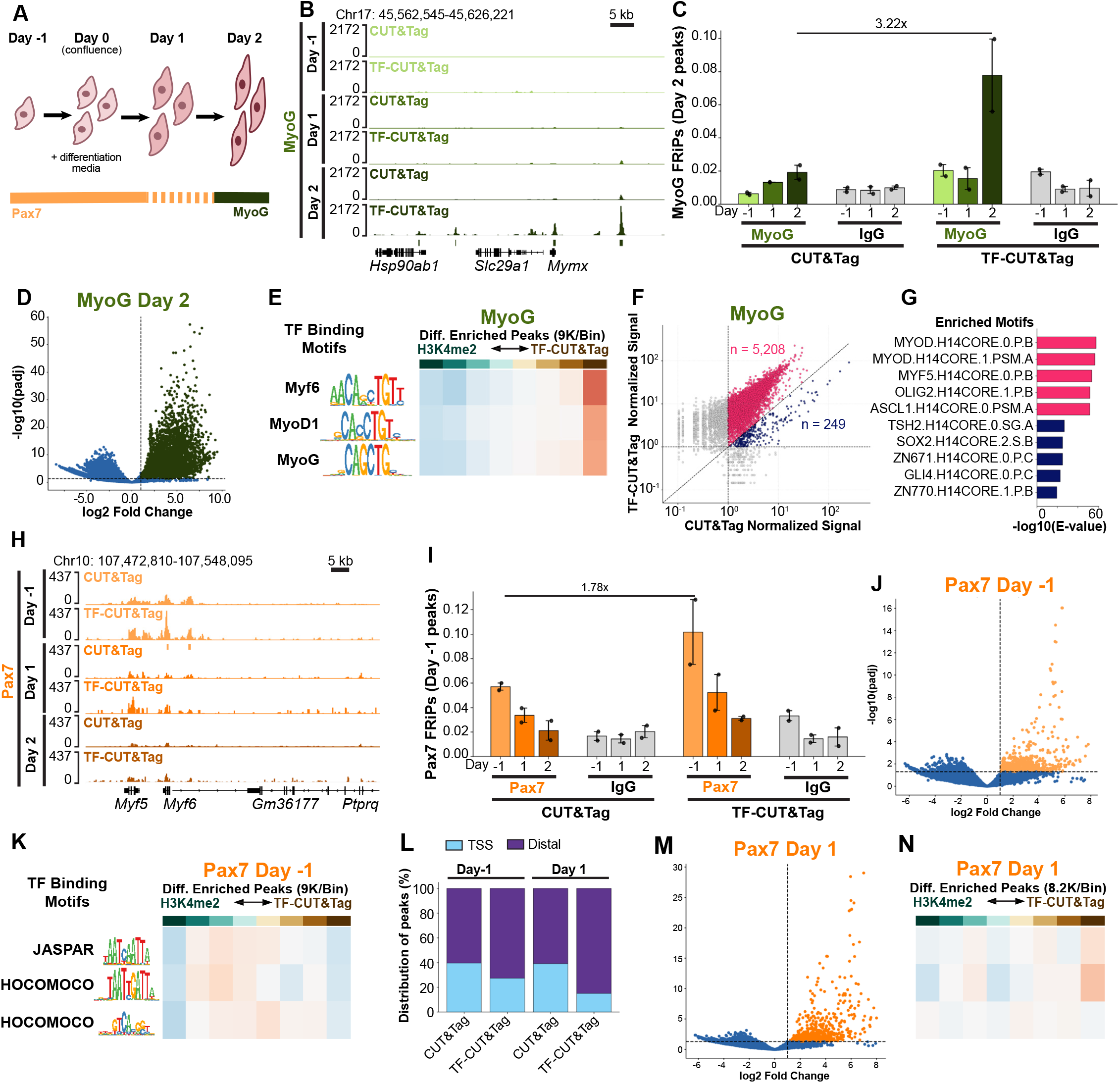
TF-CUT&Tag reveals distinct MyoG and Pax7 binding modes during myogenic differentiation. **(A)** Schematic of the C2C12 differentiation time course from proliferating myoblasts (Day −1) to differentiating cells (Days 1 and 2), with the corresponding Pax7 and MyoG expression patterns. **(B)** Genome browser tracks showing MyoG standard CUT&Tag and TF-CUT&Tag profiles over the Mymx locus at the indicated time points. **(C)** FRiP values for MyoG CUT&Tag and TF-CUT&Tag during differentiation, with matched IgG controls. **(D)** Volcano plot of DESeq2 results comparing Day 2 MyoG TF-CUT&Tag with H3K4me2 CUT&Tag. MyoG-enriched peaks (log2 fold change > 1 and adjusted P < 0.05) are shown in green, and the remaining peaks are shown in blue. monaLisa heatmap showing enrichment of myogenic E-box motifs across consecutive approximately 9,000-site bins ranked from H3K4me2-favored to MyoG TF-CUT&Tag-favored sites. **(F)** Scatterplot comparing normalized signal at merged MyoG peaks between Day 2 TF-CUT&Tag and standard CUT&Tag. Each point represents one peak; pink and navy denote TF-CUT&Tag-favored and standard-CUT&Tag-favored sites, respectively, among peaks with signal ≥1 in both conditions. **(G)** Differential AME motif enrichment in the site populations defined in (F), expressed as −log10(E-value). **(H)** Genome browser tracks showing Pax7 standard CUT&Tag and TF-CUT&Tag profiles across the Myf5/Myf6 locus during differentiation. **(I)** FRiP values for Pax7 and matched IgG controls at the indicated time points, as in (C). **(J)** Volcano plot comparing Pax7 TF-CUT&Tag at Day −1 with H3K4me2 CUT&Tag; Pax7-enriched peaks are shown in orange. **(K)** monaLisa heatmaps showing enrichment of Pax7- and Sp/KLF-associated motifs across peak bins ranked from H3K4me2-favored to Pax7 TF-CUT&Tag-favored sites at Day −1. **(L)** Stacked bar graphs showing the proportion of Pax7 peaks located in promoter/TSS-proximal or distal genomic regions for standard CUT&Tag and TF-CUT&Tag at Day −1 and Day 1. Promoter/TSS-proximal sites are shown in light blue and distal sites in purple. **(M)** Same as (J) but showing the Pax7 versus H3K4me2 CUT&Tag signal at Day 1. **(N)** Same as (K) but showing the enrichment of Pax7- and Sp/KLF-associated motifs across peak bins at Day 1.

As expected, Pax7 showed the opposite temporal pattern from MyoG, with signal progressively declining across differentiation as MyoG binding increased. For example, at the Myf5/Myf6 locus, TF-CUT&Tag recovered prominent Pax7-associated signal in the Day -1 proliferating progenitor cells that diminished by Day 2, whereas standard CUT&Tag produced weaker profiles (**Fig. 6H**). Using a merged Pax7 Day -1 reference peak set, Pax7 FRiP was highest in progenitor cells and declined during differentiation, with TF-CUT&Tag showing a 1.78-fold increase in the Pax7 FRiPs relative to standard CUT&Tag at Day -1(**Fig. 6I**). In the Day -1 proliferating progenitors where Pax7 signal was highest the canonical Pax7 motif was not preferentially enriched at the most Pax7-enriched end of the distribution (**Fig. 6J,K**). Instead Sp/KLF motifs were enriched across intermediate bins of relative Pax7 enrichment. These Pax7 sites were also disproportionately promoter-proximal and CpG-rich (**Fig. 6L**). By Day 1, however, the canonical Pax7 motif became enriched in the most Pax7-enriched bin (**Fig. 6M,N**). The emergence of canonical Pax7 motif enrichment after differentiation, together with the expected temporal decline in Pax7 signal, supports the specificity of the recovered Pax7 profiles despite the distinct sequence context observed in proliferating progenitors. Together, our results show that TF-CUT&Tag improves recovery of temporally regulated MyoG and Pax7 binding and are consistent with a differentiation-dependent shift in the sequence features associated with Pax7 binding.

## Discussion

Here, we show that attenuating Tn5 DNA engagement can improve the signal-to-background ratio of CUT&Tag for transcription factors under relatively low-salt conditions. Previous engineering of bacterial Tn5 has largely favored efficient engagement and fragmentation of accessible DNA for sequencing applications. During CUT&Tag, however, intrinsic DNA engagement can become a liability by promoting integration at accessible sites independently of antibody recruitment. The engineered variants shifted this balance in different ways, with K160A consistently improving GATA2 motif-associated recovery, and K249A performing better for profiling CTCF over sites with strong motifs and relatively low ATAC accessibility. Thus, these variants provide complementary opportunities for improving TF-CUT&Tag, and comparing TF-CUT&Tag profiles generated with the K249A and K160A variants may be particularly useful for assessing the physical chromatin environment at TF binding sites.

Many TFs engage chromatin through electrostatic interactions that can be disrupted by the stringent ionic conditions used in standard CUT&Tag. For example, IRF1 binding to its recognition sequence depends on salt concentration and identity, which influence ion release and charged protein–DNA contacts^35^. Likewise, positively charged regions flanking Sox DNA-binding domains can stabilize interactions with DNA and nucleosomes^36^. Because H3K27me3 signal was retained across the same NaCl gradient used for TF profiling, the accompanying decrease in TF-associated signal is target-dependent rather than an inevitable consequence of suppressing Tn5 activity. High-salt washes may therefore alter not only the amount of TF signal recovered, but also which classes of TF–chromatin interaction remain detectable. These effects are likely influenced by both the stability of TF binding and the ionic sensitivity of Tn5-mediated DNA engagement.

Our results indicate that differences in profiling stringency can influence not only the amount of TF-associated signal recovered, but also the types of binding sites detected. For GATA2 and MyoG, lower-stringency conditions preferentially recovered sites enriched for cognate motifs, consistent with relatively salt-sensitive, sequence-directed interactions (**Fig. 7**, Class A). For Pax7 and Nanog, differences in recovery extended across both motif-directed and less sequence-constrained sites, without a dominant shift toward the cognate motif (**Fig. 7**, Class B). Conversely, CTCF and Sox2 showed greater cognate-motif enrichment under more stringent conditions, consistent with preferential retention of strongly sequence-directed binding events (**Fig. 7**, Class C). Importantly, greater motif enrichment does not necessarily indicate a more complete TF binding profile, as it can also result from the selective loss of genuine TF– chromatin interactions.

**Figure 7.**
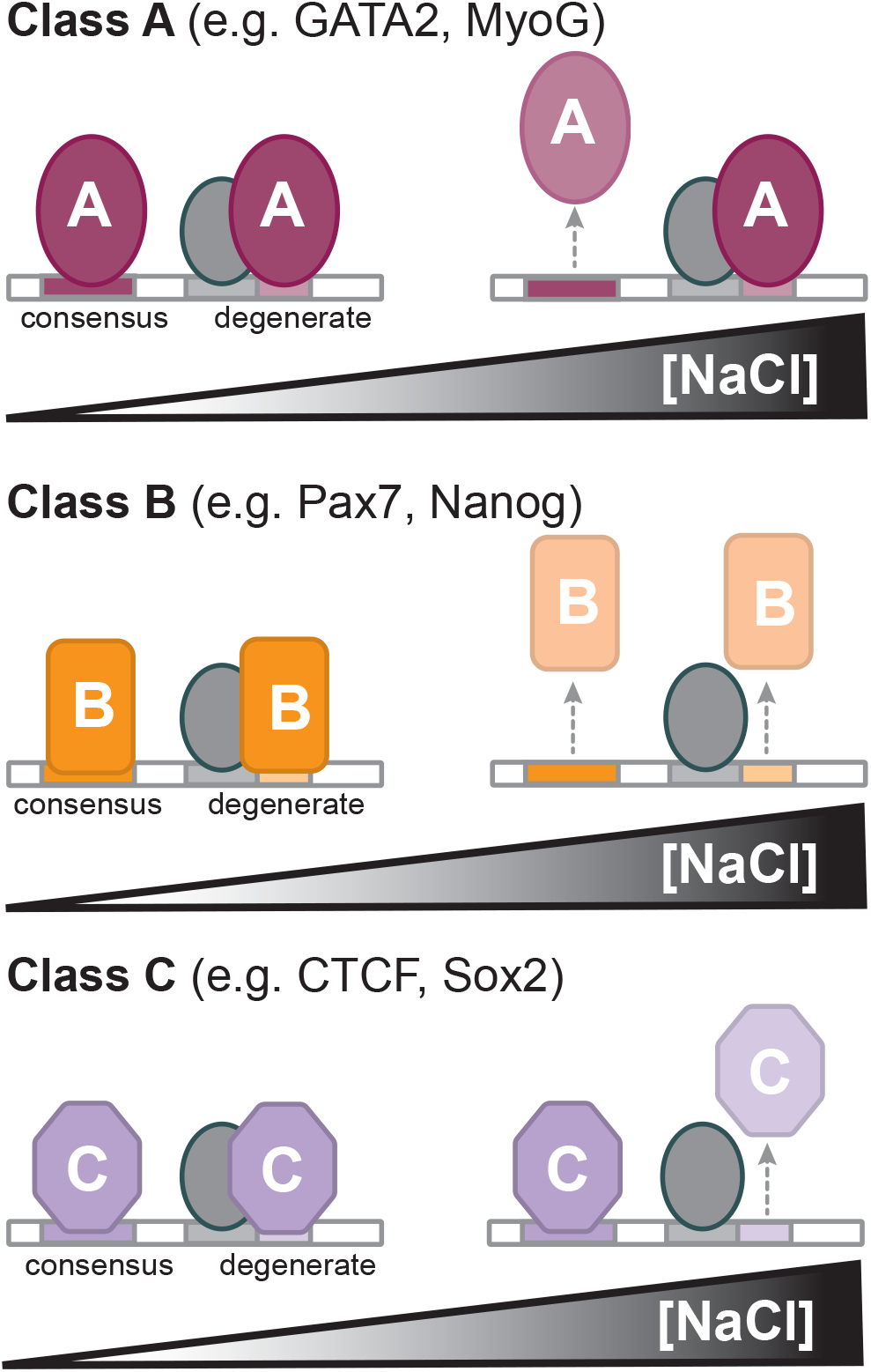
Proposed classes of transcription-factor recovery under increasing ionic stringency. Schematic illustrating three potentially overlapping patterns of TF–chromatin interactions detected by CUT&Tag as NaCl concentration increases. Colored shapes denote TF occupancy at consensus or degenerate recognition sequences, gray shapes represent possible interacting cofactors, and upward arrows indicate interactions that may be lost under more stringent conditions. Class A (e.g., GATA2 and MyoG) illustrates preferential loss of a salt-sensitive, motif-directed population. Class B (e.g., Pax7 and Nanog) illustrates sensitivity of both motif-directed and less sequence-constrained interactions. Class C (e.g., CTCF and Sox2) illustrates preferential retention of strongly sequence-directed sites despite loss of other interactions. These classes summarize assay recovery patterns and do not imply that TF-binding affinities or residence times were measured directly.

The preferential retention of motif-directed CTCF and Sox2 sites may reflect the relatively stable binding of these factors to their recognition sequences. Single-molecule studies have identified a long-lived SOX2 population at specific sites and CTCF residence times of approximately one to two minutes, exceeding those of many other TFs^37,38^. In contrast, the additional sites recovered under less stringent conditions may represent weaker sequence-specific interactions or binding supported by cofactors and protein–protein interactions. Short transcriptional activation domains can alter TF chromatin residence^9^, while intrinsically disordered regions can cooperate with transient DNA binding^7^. These mechanisms provide a plausible explanation for biologically meaningful TF binding that is difficult to predict from DNA sequence alone. Although our results reveal distinct patterns of TF recovery, they do not directly establish TF–DNA binding affinities or residence times, and some comparisons differ in both Tn5 variant and ionic conditions.

Our direct comparison of K160A and K249A suggests why the optimum may depend on chromatin context. K249A-preferred CTCF sites were less ATAC-accessible on average than K160A-preferred GATA2 sites, yet showed sharply motif-centered short-fragment enrichment (**Fig. 4**). We propose that K249A retains sufficient DNA-engagement activity to integrate within restricted windows of accessible DNA, whereas the greater attenuation of K160A can limit productive integration at some CTCF sites even after antibody recruitment. At the comparatively more accessible GATA2-associated sites, K160A may still integrate efficiently enough when tethered while strongly reducing untethered accessibility-associated tagmentation. The differing responses under DynaTag are compatible with ionic effects on DNA exposure and enzyme activity. These findings suggest that K160A may be a useful starting point for class A and B factors, whose sites can be salt-sensitive, while K249A may be advantageous for class C factors that retain stable binding but offer narrower opportunities for DNA integration.

Pax7 provides a biological example in which the relative contribution of distinct binding modes changes with cell state. In proliferative progenitor cells, Pax7-enriched sites were disproportionately promoter-proximal and CpG-rich and showed enrichment for GC-rich Sp/KLF motifs rather than the canonical Pax7 motif. As differentiation proceeded and overall Pax7 signal declined, the recovered Pax7 profile became increasingly enriched for sites containing the cognate Pax7 motif. Pax7 engages DNA through multiple binding modalities, with its paired and homeodomain DNA-binding domains supporting higher-affinity composite binding, while lower-affinity sites can be engaged primarily through the paired domain^39^. One possible explanation is a mass-action effect: as Pax7 abundance decreases during differentiation, occupancy at lower-affinity sites might decline more rapidly than occupancy at high-affinity cognate motifs. Alternatively, differentiating C2C12 cultures contain reserve cells that remain undifferentiated and enter a more quiescent state while maintaining Pax7 expression and proliferative potential^40,41^. These cells could contribute to the observed profile through changes in their representation, chromatin accessibility, interacting proteins, or Pax7 regulation. Manipulating Pax7 abundance independently of differentiation and separating differentiated and reserve-cell populations will help distinguish these possibilities. Regardless of mechanism, the coherent promoter proximity, CpG enrichment, and Sp/KLF motif composition of progenitor-associated sites suggest that they represent a biologically structured class of Pax7-associated chromatin rather than nonspecific background.

A challenge in engineering Tn5 for TF-CUT&Tag is that the desired outcome is not a simple maximum. An activity-based directed-evolution screen focused primarily on transposition efficiency would likely favor variants that strengthen the same intrinsic DNA engagement that contributes to off-target background. TF-CUT&Tag instead requires a “Goldilocks” enzyme that engages free genomic DNA less efficiently while retaining sufficient catalytic activity when tethered to an antibody-bound TF target. We therefore used a rational engineering strategy informed by biochemical analysis of a bifunctional Tn5 DNA-binding region involved in target capture^27^. Before strand transfer can occur, a preassembled Tn5 complex must capture and position target DNA. The assay used by Gradman et al. measured the combined outcome of target capture and subsequent strand transfer rather than catalytic chemistry in isolation. Because K160 is positioned away from the catalytic active site, the reduced activity of K160A in this assay was interpreted primarily as impaired target-DNA capture rather than a direct catalytic defect^27^. Although K160A showed an approximately 2.4-fold reduction in the target-capture/strand-transfer assay, our results suggest that this attenuation becomes advantageous when the enzyme is tethered. For an untethered Tn5 molecule encountering accessible genomic DNA by diffusion, reducing the probability of productive target capture should decrease the frequency with which a random encounter proceeds to insertion. Antibody tethering, in contrast, maintains a high local concentration of Tn5 adjacent to the target chromatin, and provides repeated opportunities for productive capture even when individual Tn5–DNA interactions are weak. A modest reduction in intrinsic target-capture efficiency may therefore preferentially penalize untethered tagmentation while providing multiple opportunities for tagmention at antibody-bound sites. However, reduced activity alone was not sufficient to improve TF-CUT&Tag. R104A reduced activity *in vitro* by approximately 2.6-fold but did not improve TF specificity as measured by motif enrichment, whereas H213A reduced activity by approximately 5.3-fold and produced a modest specificity improvement that remained weaker than K160A^27^. Other substitutions more severely compromised productive CUT&Tag activity. Optimal performance therefore likely depends on the balance among target capture, transposome stability, catalytic competence, and the ability of tethering to compensate for reduced intrinsic DNA engagement. K160A appears to bind a favorable region of this functional landscape. Direct measurements of target-DNA association and dissociation, equilibrium affinity, target positioning, and catalytic strand transfer could define the molecular basis of this behavior and guide a second generation of Tn5 variants optimized for tethered chromatin profiling.

There is precedent for tuning rather than maximizing Tn5 activity for specialized genomic applications. The R27S mutation was introduced accidentally during PCR amplification and cloning and was subsequently found to make DNA fragmentation strongly dependent on the ratio of Tn5 to input DNA, allowing fragment size to be controlled by enzyme concentration^42^. This property was later exploited in long-read applications in which controlled tagmentation and preservation of long native DNA molecules are advantageous^43^. R27 contacts the mosaic-end-containing adapter DNA loaded into Tn5, whereas K160 is implicated in capture of the incoming genomic target DNA. Thus, R27S and our K249A and K160A alter different steps of the tagmentation reaction, but both illustrate how weakening or tuning an otherwise advantageous Tn5–DNA interaction can improve the enzyme for a specialized genomic application.

An advantage of engineered TF-CUT&Tag is that it retains the basic CUT&Tag workflow and can build directly on existing high-throughput, single-cell, and multifactorial implementations. Automated approaches such as AutoCUT&Tag could screen K160A, K249A to identify the best Tn5 variant-to-TF pairings and probe the physical chromatin at sites bound by various TFs^18^. Extensions to multifactorial and single-cell methods such as MulTI-Tag and CoCUT&Tag could then assess these optimized profiles in heterogeneous tissues and transient developmental states^21,44^. More broadly, our results indicate that Tn5 can be engineered to favor tethered over untethered integration, providing a route to continued optimization of TF-CUT&Tag fidelity and a unified enzyme-tethering method for profiling TF binding sites.

## Resource availability

### Lead Contact

Questions related to this study and requests for resources and reagents should be directed to the lead contact, Derek Janssens, and will be fulfilled in a timely manner.

### Materials Availability

Materials generated in this study will be available upon reasonable request. The completed K160A construct (Addgene plasmid # 264171), is available through Addgene.

### Data and Code Availability

The CUT&Tag, TF-CUT&Tag, and DynaTag primary sequencing data are available through the Gene Expression Omnibus under the accession number: GSE351354.

## Acknowledgments

We thank Drs. Steve Henikoff and Terri Bryson for kindly sharing the K562 cells used in this study, and Dr. Tristan Kempston and the VAI transgenics core for sharing the B6 mouse ESCs used for profiling the pluripotency TFs. We thank Drs. Weifang Wu, Darrell Chandler, Sophia Chaudhry, Tim Triche Jr., Johnathan Licht, as well as all members of the Janssens lab and the rotation students from the VAI graduate class of 2025 for helpful discussion and feedback on the manuscript. We thank all members of the Van Andel Institute Genomics Core (RRID:SCR_022913) for technical assistance with sequencing and data processing. We are also grateful for Jack Bidwell for creative direction on figure panels, and Drs Martin Burg, Matthew Christans, Sok-Kean Khoo, and Mark Staves for their support and encouragement throughout this project. This work was supported by grants from the National Institutes of Health NCI K22 1K22CA272912-01A1 (D.H.J.), a Scholar Award from the American Society for Hematology (D.H.J), and the Meijer Foundation (L.M.P). Work by S.M.N. (Sara M. Nowinski) and O.A.P. (Olivia A. Pentecost) was supported by the National Institute of Arthritis and Musculoskeletal and Skin Diseases (NIAMS), National Institutes of Health, under award R01AR085095.

## Author Contributions

D.H.J. and E.B. conceived the study. L.M.P, E.B., H.J.A, O.A.P, and D.H.J. performed the experiments; L.M.P, E.B. and D.H.J. performed data analysis and assembled figures.

L.M.P and D.H.J. wrote the manuscript; L.M.P, S.M.N, and

D.H.J reviewed and edited the manuscript, and all authors approved the manuscript.

## Methods

### Cloning plasmids for bacterial expression of TF-CUT&Tag proteins

The backbone plasmid used for all cloning was the 3XFlag-pA-Tn5-Fl, which was a gift from Steven Henikoff (Addgene plasmid # 124601 ; RRID:Addgene_124601). The QuikChange Lightning Multi Site-Directed Mutagenesis Kit (Agilent, 210515) was used to create plasmids with each of the single amino acid substitutions characterized herein, following the manufacturer’s protocol except that DpnI digestion was performed with 3 µL of enzyme overnight rather than the recommended amount and time.

### Protein Purification

All the pA-Tn5 variants tested for TF-CUT&Tag were expressed with a C-terminus fusion to an intein and chitin binding domain (CBD). The CBD specifically binds chitin resin, and addition of a reducing agent (Dithiothreitol, DTT) causes the intein to cleave and release the target protein into solution. Each variant pA-Tn5 protein was obtained using the same protein expression and purification workflow. Each construct was transformed into T7 Express lysY/ Iq Competent *E. coli* (NEB #C3013I) following the manufacturer’s protocol and plated for selective overnight growth. A single colony was used to inoculate 5 mL of LB media containing 100 µg/ml carbenicillin. The starter culture was incubated overnight at a temperature of 37°C while shaking at 220 RPM. The 5 mL starter culture was then used to inoculate 345 mL TB media containing 100 µg/ml carbenicillin in a 1L baffled flask. The 350 mL culture was incubated at a temperature of 37°C while shaking at 170 RPM until the OD600 reached 0.75-1.00 (approximately two hours). After reaching this density, the culture was placed in a cold room for one hour to cool then promptly induced via the addition of IPTG to a final concentration of 0.25 mM. The induced culture was incubated overnight at a temperature of 18°C while shaking at 170 RPM. The bacterial culture was pelleted by centrifugation for 10 minutes at 9,500 × *g*. All centrifugations were performed at a temperature of 4°C. The bacterial pellet was flash frozen using liquid nitrogen and stored at a temperature of -80°C. To proceed, the pellet was thawed on ice in a 4°C cold room. A stir plate was then used to resuspend the pellet in 35 mL chilled HEGX (0.02 M HEPES-KOH at pH 7.2, 1 M NaCl, 0.001 M EDTA, 10% glycerol, 0.2% Triton X-100, and fresh protease inhibitor added immediately before use (Sigma #5056489001)). The resuspension was sonicated in ice-nested beakers with a microtip set to 40% amplitude for 12 rounds: 3 seconds on, 3 seconds off, for 40 seconds total as detailed by Bryson & Henikoff (https://dx.doi.org/10.17504/protocols.io.8yrhxv6). The lysate was then centrifuged at 29,000 × *g* for 30 minutes. A chitin resin column was prepared in a 4°C cold room by gently inverting the resin (NEB #S6651) to resuspend and adding 2.5 mL resin to an Econo-Pac Chromatography Column (Bio-Rad #7321010). With the column set upright in a column rack, the resin was allowed to settle and then washed twice with 20 mL HEGX Buffer. Once the HEGX Buffer had run through the column, the bottom of the column was capped, and 20 mL of the soluble lysate (i.e., supernatant) was added. The column was capped on the top, parafilmed, and left nutating overnight at 4°C. The next day, the column was returned to the rack, and the unbound fraction was drained into a collection tube. After washing the column twice with 20 mL HEGX Buffer and recapping the bottom, 6 mL HEGX Buffer containing 100 mM DTT and fresh protease inhibitor was added slowly to the column. The column was capped on the top, parafilmed, and incubated for 48 hours at 4°C. The DTT-cleaved fraction was then drained into a collection tube. The collected sample was loaded in a dialysis cassette (Life Tech #66810) following product literature and dialyzed overnight in Dialysis Buffer (0.1M HEPES-KOH at pH 7.2, 0.2 M NaCl, 0.2 mM EDTA, 0.2% Triton X-100, 20% glycerol, with DTT added to a final concentration of 1.7 mM immediately before use) with a buffer change after two hours. The dialyzed sample was recovered from the cassette and centrifuged at 3000 × *g* for two minutes to remove precipitated protein. The sample was then held on ice at 4°C until the protein was verified on an SDS-PAGE gel. After verification, the Tn5 proteins were loaded with adapter oligos and stored, as is further described. The solution containing Tn5 proteins can also be snap frozen and stored at -80°C for loading up to one year later.

### Adapter annealing and Tn5 loading

Purified Tn5 mutant proteins were loaded with Tn5MEDS-A and Tn5MEDS-B oligonucleotides pre-annealed to the universal ME-reverse oligonucleotide (Picelli et al., 2014). The oligonucleotides were ordered from IDT with standard desalting and resuspended to 200 µM in Annealing Buffer (10 mM Tris-HCl pH 8.0, 50 mM NaCl, 1 mM EDTA). An equal volume of 200 µM universal ME-reverse oligonucleotide was mixed separately with Tn5MEDS-A and Tn5MEDS-B, and each mixture was annealed in a thermocycler with a 105°C heated lid. The program began with a 2-minute incubation at 95°C, decreased in 5°C increments with a 5-minute incubation at each step to 25°C, and then held at 8°C. Pre-annealed oligonucleotides were used immediately or stored at −20°C.

Pre-annealed Tn5MEDS-A/ME-reverse and Tn5MEDS-B/ME-reverse were mixed at equimolar concentrations and added to Tn5 mutant protein at a volume ratio of 1:1:12.5. Protein concentration was not determined before loading; oligonucleotides were added in excess. The solution was vortexed, and loading proceeded for 1.5 hours at room temperature. An equal volume of 100% glycerol was then added (50% final), and loaded proteins were stored at −20°C.

### Agarose gel analysis of loading

Loading was assessed by comparing the amount of oligonucleotide bound to mutant versus parental Tn5 loaded in parallel under the same conditions (**Supplementary Figure 2B**). A 4% agarose gel was prepared in fresh 1× TBE. Loaded protein samples (25 µL) were mixed with 5 µL 6× TrackIt Cyan/Yellow loading buffer (Invitrogen 10482035). Oligonucleotide controls were (A) pre-annealed Tn5MEDS-A/ME-reverse, (B) pre-annealed Tn5MEDS-B/ ME-reverse, and (C) an equimolar mixture of both. Controls were diluted into 1× TBE (1 µL of A or B, or 2 µL of C, to 25 µL final volume) to approximate the oligonucleotide amount in the protein lanes, then mixed with 5 µL 6× TrackIt. The ladder (Invitrogen 10597012) was mixed with 6× TrackIt and nanopure water at 1:1:4. Ladder (4.8 µL) and 7.5 µL of each control and protein sample were loaded, and the gel was run at 60 V for 3.5 hours. The gel was stained with 1× SYBR Gold (Invitrogen S11494) for 40 minutes at room temperature with rotation and imaged on a Bio-Rad ChemiDoc MP Imaging System using Image Lab Touch Software (version 3.0.1.14) with the SYBR Gold preset. Unadjusted images were exported as 16-bit TIFF files for analysis.

### SDS-PAGE quantification of Tn5 abundance

Unloaded Tn5 proteins (12 µL) were mixed with 3 µL 5× Loading Buffer (stock composition: 60 mM Tris-HCl pH 6.8, 2% SDS, 10% glycerol, 0.025% bromophenol blue, 75 mM β-mercaptoethanol), boiled for 5 minutes, and the total 15 µL was resolved on a precast 4–20% polyacrylamide gel (BioRad 4561093). Each gel included a BSA standard at 125 ng/µL (prepared at the same volume as samples; Invitrogen 23208) from a pre-diluted standard set. Gels were stained with Coomassie and imaged on the ChemiDoc MP using the Coomassie preset (Supp. 2A). The unadjusted image was exported as a 16-bit TIFF file for further analysis.

### Densitometry and calculation of loading efficiency

Gel bands were quantified in ImageJ (NIH, version 1.54g). A single fixed-size rectangular ROI encompassing a full band was used for every band. For each band, ROI integrated density (IntDen) was measured, and background was subtracted using an identically sized ROI placed in an empty region of the same gel: corrected IntDen = sample IntDen – blank IntDen. The same procedure was applied to the SDS-PAGE gels described above. Protein concentration (ng/µL) was calculated using the SDS-PAGE gels as (sample corrected IntDen / BSA corrected IntDen) × 125 ng/µL, using the single BSA standard from the same gel. On the agarose gel, oligonucleotide-loaded Tn5 only migrates a short distance into the gel, and the remaining (unbound) oligonucleotide migrates further down. Only the retained signal near the well was quantified. The amount of oligonucleotide bound by mutant Tn5s was assessed relative to the amount bound by the parental Tn5 as sample corrected IntDen / parental pA-Tn5 corrected IntDen. Variation in relative oligonucleotide loading is only partially explained by Tn5 protein concentration (y = 0.0024x + 0.614; R^2^ = 0.3829; Supp. 2C). Values are from a single loading experiment, and the plot was prepared in Microsoft Excel.

### Structural modeling of pA-Tn5

A model of the pA-Tn5 dimer bound to adapters and a random DNA sequence was generated with AlphaFold Server (AlphaFold 3; Abramson et al., 2024) using the parental sequence. The top-ranked model was used for analysis and visualized in PyMOL (v3.1.3, Schrödinger, LLC). Model confidence was high (ipTM = 0.88, pTM = 0.90).

### Antibodies

Of the rabbit and mouse monoclonal antibodies we tested, we found the following antibodies performed the best in CUT&Tag and TF-CUT&Tag assays: rabbit anti-H3K27me3 (Cell Signaling Technologies; cat. no. 9733, rabbit anti-H3K4me2 (Cell Signaling Technologies; cat. no. 9725), rabbit anti-H3K4me2 (Sigma Aldrich; cat. no. 07-030), mouse anti-H3K4me2 (Active Motif; cat. no. 39679). For standard CUT&Tag and TF-CUT&Tag assays using a rabbit primary antibody, the secondary guinea pig anti-rabbit IgG antibody (Antibodies Online, cat. no. ABIN101961) served as an adaptor to amplify the number of tethered pA-Tn5 complexes. To amplify the signal from mouse primary antibodies, we used the secondary rabbit anti-mouse IgG antibody (Abcam; cat. no. ab46540). For CUT&Tag, TF-CUT&Tag, and DynaTag in K562 cells, we used a polyclonal GATA2 (Life Technologies; cat. no. PA1-100), and rabbit monoclonal anti-CTCF (Abcam; cat. no. ab128873). For CUT&Tag and TF-CUT&Tag in B6 ESCs, we used rabbit monoclonal anti-Oct4 (Abcam; cat. no. ab109183), rabbit monoclonal Sox2 (Cell Signaling Technologies; cat. no. 23064), rabbit monoclonal Nanog (Cell Signaling Technologies; cat. no. 8822), rabbit monoclonal anti-c-Myc (Abcam; cat. no. ab32072). For CUT&Tag and TF-CUT&Tag in C2C12 myoblasts, we used monoclonal mouse Pax7 (DSHB; cat. no. PAX7;AB_528428) and mouse monoclonal F5D (DSHB; cat. no. F5D;AB_2146602).

### Testing TF-CUT&Tag on a Salt Gradient

TF-CUT&Tag was performed using a direct-to-PCR protocol like the one previously described^16^. Briefly, whole cells were spun down at 600 × g and brought up in 900 μL Wash Buffer containing Triton-X 100 for permeabilization (48 mL ultrapure water, 1 mL 1M HEPES pH 7.5 [20 mM], 1.5 mL 5M NaCl [150 mM], 12.5 μL 2M Spermidine [0.5 mM], 250 μL of 10% Triton-X-100 [0.05%],1 Roche Complete EDTA-free tablet). ConA magnetic beads (Bangs, cat. no. BP531) were prepared by washing 5 μL per sample, 2 times with 500 μL Binding Buffer (400 μL of 1M HEPES-KOH pH 7.9 [20 mM], 200 μL of 1M KCl [10 mM] 20 μL of 1M CaCl2 [1 mM], and 20 μL of 1M MnCl2 [1 mM], brought up to 20 mL with ultrapure water). ConA Beads were resuspended in 100 μL of Binding Buffer and were added to the whole cell suspensions in three 33 μL increments, the sample was mixed by inversion after each addition. Bead binding was allowed to proceed for 10 min at room temperature, and the supernatant was removed on a magnet stand. Then cells were washed once with 500 μL of Wash Buffer and brought up in 15 μL of Antibody Buffer (1 mL Wash Buffer, 4 μL 0.5M EDTA [2 mM]) per TF-CUT&Tag reaction/replicate. 1.5 μL of the rabbit or mouse primary antibody (e.g, rabbit-anti H3K27me3; Cell Signaling Technologies Cat No. 9733S) was added, and antibody binding was allowed to proceed for one hour at room temperature. Cells were then washed twice with 100 μL of Wash Buffer and brought up in 25 μL of Wash Buffer and 1.5 μL of secondary antibody (either guinea pig anti-rabbit IgG antibody; Antibodies Online Cat No. ABIN101961 or rabbit anti-mouse IgG antibody; Abcam Cat No. Ab46540) was added to each tube and incubation was allowed to proceed at 4°C for one hour. Cells were then washed twice with 100 μL of Wash Buffer and brought up in 15 μL of a NaCl-Wash Buffer at the NaCl molarity being tested. 300-Wash Buffer (48 mL ultrapure water, 1 mL 1M HEPES pH 7.5 [20 mM], 3 mL 5M NaCl [300 mM], 12.5 μL 2M Spermidine [0.5 mM], 250 μL of 10% Triton-X-100 [0.05%], 1 Roche Complete EDTA-free tablet), 250-Wash Buffer (same as 300-Wash Buffer except 2.5 mL 5M NaCl [250 mM]), 200-Wash Buffer (same as 300-Wash Buffer except 2.0 mL 5M NaCl [200 mM]), 150-Wash Buffer (same as 300-Wash Buffer except 1.5 mL 5M NaCl [150 mM]). Then 1.5 μL of a variant pA-Tn5 purified and loaded according to standard protocols (dx.doi.org/10.17504/protocols.io.8yrhxv6) was added and incubation was allowed to proceed at 4°C for one hour. Cells were then washed twice with 100 μL of the appropriate NaCl-Wash Buffer (same as previous step) and brought up in 50 μL of the appropriate NaCl-Tagmentation Buffer (e.g., 300-Tagmentation Buffer: 990 μL 300-Wash Buffer + 10 μL 1M MgCl2 [10 mM]), and tagmentation was carried out at 37°C overnight in a water bath. Cells were then washed one time with TAPS Buffer (500 μL 1M TAPS pH 8.5 [10 mM] in 49.5 mL ultrapure water), and then brought up in 5 μL SDS Release Buffer (220 μL 10 mM TAPS, 25 μL 1% SDS [0.1% final], 5 μL Thermolabile Proteinase K). Samples were incubated on the thermocycler with a heated lid at 37°C for 1 hour then 58°C for one hour. The SDS was then quenched by addition of 4 μL of 1.5% Triton-X-100 (75 μL of 10% TritonX-100 solution in 425 μL of ultrapure water). Each sample was then diluted by addition of 24 μL of ultrapure water, followed by 18 μL of PCR “Master Mix” (for each sample: 10 μL 5 X HiFi buffer, 5.5 μL ultrapure water, 1.5 μL 10 mM dNTP, 1 μL KAPA HiFi Polymerase). 2 μL of a barcoded Forward primer (10 μM) and 2 μL of a barcoded Reverse primer (10 μM) were added to each sample, ensuring that each sample had at least one distinct barcode. Samples were mixed by vortexing, then placed in the Thermocycler and run on the following protocol with a heated lid at 105°C: 58°C for 5 min (gap fill), 72°C for 5 min, 98°C for 45 seconds, followed by 14 cycles of (98°C for 15 seconds, 60°C for 10 seconds, 72°C for 1 seconds), ending with 72°C for 1 minute and hold at 12°C. Samples were then purified using a standard Ampure bead cleanup with a 1.3 X size selection and then run on a Tape Station and pooled at equimolar concentrations for sequencing on the Element Aviti Platform.

### DynaTag

DynaTag was performed as previously described^26^, with the exceptions that cells were bound to ConA beads throughout, and a direct-to-PCR protocol was used during library prep^16^. Briefly, whole cells were spun down at 600 × g and brought up in 500 μL DynaTag Wash Buffer containing Triton-X 100 for permeabilization (43 mL ultrapure water, 1 mL 1M HEPES pH 7.5 [20 mM], 5.5 mL 1M KCl [110 mM], 100 μL 5M NaCl [10 mM], 50 μL 1M MgCl_2_ [1 mM], 0.5 g BSA [1% w/v], 12.5 μL 2M Spermidine [0.5 mM], 250 μL of 10% Triton-X-100 [0.05%], 1 Roche Complete EDTA-free tablet). ConA magnetic beads (Bangs, cat. no. BP531) were prepared by washing 5 μL per sample, 2 times with 500 μL Binding Buffer (400 μL of 1M HEPES-KOH pH 7.9 [20 mM], 200 μL of 1M KCl [10 mM] 20 μL of 1M CaCl_2_ [1 mM], and 20 μL of 1M MnCl_2_ [1 mM], brought up to 20 mL with ultrapure water). ConA Beads were resuspended in 100 μL of Binding Buffer and were added to the whole cell suspensions in three 33 μL increments, and the sample was mixed by inversion after each addition. Bead binding was allowed to proceed for 10 min at room temperature, and the supernatant was removed on a magnet stand. Then cells were washed once with 500 μL of DynaTag Wash Buffer and brought up in 15 μL of DynaTag Antibody Buffer (1 mL DynaTag Wash Buffer, 4 μL 0.5M EDTA [2 mM]) per DynaTag reaction/replicate. 1.5 μL of the rabbit or mouse primary antibody (e.g, rabbit-anti H3K27me3; Cell Signaling Technologies Cat No. 9733S) was added, and antibody binding was allowed to proceed overnight at 4°C. The next day, cells were washed twice with 100 μL of DynaTag Wash Buffer and brought up in 25 μL of DynaTag Wash Buffer and 1.5 μL of secondary antibody (either guinea pig anti-rabbit IgG antibody; Antibodies Online Cat No. ABIN101961 or rabbit anti-mouse IgG antibody; Abcam Cat No. Ab46540) was added to each tube and incubation was allowed to proceed at 4°C for one hour. Cells were then washed twice with 100 μL of DynaTag Wash Buffer and brought up in 15 μL of DynaTag Wash Buffer. 1.5 μL of pA-Tn5 purified and loaded according to standard protocols (dx.doi.org/10.17504/protocols.io.8yrhxv6) was added and incubation was allowed to proceed at 4°C for one hour. Cells were then washed twice with 100 μL of DynaTag Wash Buffer and brought up in 50 μL of DynaTag Tagmentation Buffer (990 μL DynaTag Wash Buffer + 10 μL 1M MgCl2 [10 mM]), and tagmentation was carried out at 37°C for 1 hour on a thermocycler with a heated lid. Cells were then washed one time with TAPS Buffer (500 μL 1M TAPS pH 8.5 [10 mM] in 49.5 mL ultrapure water), and then brought up in 5 μL SDS Release Buffer (220 μL 10 mM TAPS, 25 μL 1% SDS [0.1% final], 5 μL Thermolabile Proteinase K). Samples were incubated on the thermocycler with a heated lid at 37°C for 1 hour then 58°C for one hour. The SDS was then quenched by addition of 4 μL of 1.5% Triton-X-100 (75 μL of 10% TritonX-100 solution in 425 μL of ultrapure water). Each sample was then diluted by addition of 24 μL of ultrapure water, followed by 18 μL of PCR “Master Mix” (for each sample: 10 μL 5 X HiFi buffer, 5.5 μL ultrapure water, 1.5 μL 10 mM dNTP, 1 μL KAPA HiFi Polymerase). Then 2 μL of a barcoded Forward primer (10 μM) and 2 μL of a barcoded Reverse primer (10 μM) were added to each sample, ensuring that each sample had at least one distinct barcode. Samples were mixed by vortexing, then placed in the Thermocycler and run the following protocol with a heated lid at 105°C: 58°C for 5 min (gap fill), 72°C for 5 min, 98°C for 45 seconds, followed by 14 cycles of (98°C for 15 seconds, 60°C for 10 seconds, 72°C for 1 seconds), ending with 72°C for 1 minute and hold at 12°C. Samples were then purified using a standard Ampure bead cleanup with a 1.3 X size selection and then run on a Tape Station and pooled at equimolar concentrations for sequencing on the Element Aviti Platform.

### Myoblast cell culture

C2C12 immortalized mouse skeletal myoblasts (ATCC CRL-1772) were maintained in DMEM with 4.5 g/L glucose, glutamine, and sodium pyruvate (Corning, 10–013-CV) with 10% FBS (Sigma, F0926) at 37°C in a humidified incubator containing 5% CO_2_.

### Myoblast differentiation assays

C2C12 myoblasts were seeded and allowed to grow to 95% confluency in DMEM (Corning, 10–013-CV) with 10% FBS (Sigma, F0926). Once at 95% confluency, cells received a PBS rinse and were switched to a differentiation media consisting of DMEM (Corning, 10–013-CV) with 2% horse serum (Thermo, 16050122) and received a differentiation media change every 24 hours. Samples were harvested using Accutase (Thermo, A11105-01) 24 or 48 hours after differentiation media was added.

### Quantifying H3K27me3 on-target and off-target signal for pA-Tn5 mutants across a salt gradient

Replicate BED files were combined (zcat) and sorted using BEDTools (v2.30.0) for both H3K27me3 CUT&Tag and ATAC-seq datasets. The combined H3K27me3 (300 mM, control) BED file was converted to a bedGraph intermediate (BEDtoBIGWIG) and peaks were called using SEACR in “non” mode with an FDR threshold of 0.1. The combined ATAC-seq BED file was downsampled to 0.1 using BEDTools, and peaks were called using MACS3 as described above, using the hg38 genome assembly, an extension size of 200 bp, and no p-value/q-value cutoff. An accessibility-associated peak set was defined by intersecting the ATAC-seq and H3K27me3 peak files (bedtools intersect) to retain only ATAC-seq peaks not overlapping H3K27me3 signal.

Normalized signal over the H3K27me3 and accessibility-associated peak sets was calculated for each sample by computing coverage per peak, normalized by peak size and by total sequencing depth: Signal_norm = (overlap bp / peak size) × (1×10□ / total bp) + 1. Peaks with no overlapping reads were retained in this calculation with a coverage value of zero rather than being dropped. Normalized signal was plotted on a log-scale y-axis as boxplots grouped by buffer or mutant condition and peak set, and compared between conditions (e.g., mutant vs. control Tn5, or one NaCl concentration vs. another) using a two-sided Welch’s t-test (unequal variances; scipy.stats. ttest_ind, equal_var=False), with results reported as the median difference in normalized signal (mutant − control, or lower-salt − higher-salt, as appropriate) and corresponding p-values.

### Differential analysis of CUT&Tag/TF-CUT&Tag for transcription factors against ATAC-seq or H3K4me2

Replicate BED files for each condition were combined (zcat) and sorted using BEDTools (v2.30.0), and peaks were called using MACS3 (macs3 callpeak -t <BED> -f BED -g hs --nomodel --extsize 200). A fixed-width peak file of 200 bp summits was also generated for each condition. To compare two conditions, their 200 bp summit files were merged using a sequential nearest-neighbor approach, retaining the higher-scoring summit at any overlapping positions. For comparisons in K562 cells, CUT&Tag/TF-CUT&Tag peaks were merged with ATAC-seq peaks in this way; for C2C12 and mESC experiments, TF-CUT&Tag peaks were merged with H3K4me2 CUT&Tag peaks.

Read counts per peak were generated for each sample by intersecting its BED file against the merged peak file, producing a count matrix across all samples being compared. Differential analysis was carried out on this count matrix using DESeq2 in R, and results were imported into Jupyter and plotted as volcano plots (log2 fold change on the x-axis, −log10(adjusted p-value) on the y-axis). Peaks with a fold change greater than 2 and an adjusted p-value less than 0.05 were exported as a BED file, filtered for canonical chromosomes, and labeled “enriched” for the comparison condition (e.g., the TF); all remaining peaks were similarly exported and labeled “not enriched.”

“Enriched” and “not enriched” peak sets were converted to FASTA files using BEDTools getfasta with the appropriate reference genome (hg38 for K562; mm10 for C2C12 and mESC experiments) and analyzed for differential motif enrichment using AME with the HOCOMOCO v14 core motif database (with hg38-exclusive motifs removed for mm10 analyses), average odds scoring, and a Fisher’s exact test for motif enrichment as parameters. AME was run in both directions for each comparison with “enriched” peaks as input against “not enriched” as background, and vice versa to identify motifs enriched in each direction. For mESC comparisons, the resulting enrichment values (−log10(E-value)) for the expected Yamanaka factor associated motifs were plotted as bar charts.

### Head-to-head comparisons of CUT&Tag conditions

#### Signal boxplots

GATA2 and CTCF signal was quantified over the “enriched” (TF) and “not enriched” (ATAC-seq) peak sets exported from the DESeq2 analysis described above, using the same normalization, plotting, and statistical approach detailed in the H3K27me3 section.

#### Motif enrichment analysis

For each transcription factor, a master peak set was generated by merging 200 bp summit peaks across replicates from every relevant Tn5 condition, using the same summit-merging approach described above. Any comparisons including DynaTag had DynaTag replicates included in the merged peakset. For K562 experiments, all replicates from every Tn5 mutant across all salt buffer conditions were included; for C2C12 and mESC experiments, only TF-CUT&Tag (K160A, 200 mM NaCl) and standard CUT&Tag (control, 300 mM NaCl) were included. To compare signal between two conditions (e.g., mutant vs. control Tn5, DynaTag vs CUT&Tag) over this merged peak set, reads from each replicate BED file were counted per peak using BEDTools and normalized to reads per million (total reads × 1×10╁), then averaged across replicates. Normalized signal from the two conditions was plotted against one another as scatter plots, with the reference condition on the x-axis and the comparison condition on the y-axis.

Condition favored peaks were defined using three thresholds on these scatter plots: a minimum normalized signal of 1 on each axis, and the diagonal (y = x). Peaks meeting the signal threshold were classified as “comparison-favored” if they fell above the diagonal, or “control-favored” if they fell below it, and highlighted in pink and navy, respectively. These peak sets were converted to FASTA files and analyzed for differential motif enrichment using AME, run in both directions, exactly as described in the DESeq2-based motif analysis above. For K562 comparisons, results were displayed as heatmaps of signed −log10(E-value), with pink indicating enrichment of the expected TF motif (GATA2 or CTCF) in the comparison condition and navy indicating enrichment in the control condition. For C2C12 comparisons, results were displayed as bar charts of the top five enriched motifs in each direction (comparison in pink, control in navy).

#### FRiP analysis

Fraction of reads in peaks (FRiP) was calculated for each transcription factor under both standard CUT&Tag and TF-CUT&Tag (K160A), comparing each antibody condition to its matched IgG control. In each case, FRiP was calculated over a single merged peak set built by combining peaks from all relevant CUT&Tag and TF-CUT&Tag conditions (with and without antibody) for that factor, rather than using a peak set unique to each individual condition.

For the C2C12 myogenesis time course (Pax7, MyoG), this merged peak set was built from the differentiation stage at which each factor’s expression is expected — Dp2 (late differentiation/myotubes) for MyoG, and Dn1 (proliferating myoblasts) for Pax7 — combining peaks called from both antibody and IgG data, and from both standard CUT&Tag and TF-CUT&Tag, at that stage. This single peak set was then used to calculate FRiP across all three differentiation stages (Dn1, Dp1, Dp2) for that factor, for each antibody condition (Dn1 + antibody, Dp1 + antibody, Dp2 + antibody) compared to IgG at the matched time point. Fold-change in FRiP between CUT&Tag and TF-CUT&Tag is reported for the relevant comparison (e.g., 3.22-fold for MyoG at Dp2, 1.78-fold for Pax7 at Dn1).

For the mESC Yamanaka factor analysis (Oct4, Sox2, Nanog), the merged peak set for each factor was built from four conditions: standard CUT&Tag with the TF antibody, standard CUT&Tag with IgG, TF-CUT&Tag with the TF antibody, and TF-CUT&Tag with IgG.

### Motif enrichment analysis of DESeq results using monaLisa

Sites were the merged peak sets used for each DESeq2 comparison: 200 bp MACS3 summits from each condition were merged by retaining the higher-scoring summit at overlapping positions. TF-CUT&Tag and CUT&Tag peaks were used for GATA2. TF-CUT&Tag with H3K4me2 CUT&Tag peaks were used for Oct4, Sox2, Nanog, cMyc, MyoG, and Pax7.

Reads from each replicate were counted per site with bedtools intersect against the merged peak set. Differential signal between TF-CUT&Tag (K160A, 200 mM NaCl) and H3K4me2 CUT&Tag (for C2C12 and mESC experiments) or GATA2 TF-CUT&Tag (K160A, 200 mM NaCl) and GATA2 CUT&Tag (for GATA2 experiments) was tested with DESeq2 (v1.52.0) using a negative binomial model with the design ∼ treatment + replicate [n = 2 replicates per condition], default median-of-ratios normalization and Wald tests, with Benjamini–Hochberg adjustment. Log2 fold changes were not shrunk. Positive values indicate higher signal in TF-CUT&Tag.

Binned motif enrichment analysis was performed with monaLisa (v1.18.0) in R (v4.6.1, Bioconductor v3.23) on DESeq2 results comparing TF-CUT&Tag (K160A) to a reference condition for each factor. For the Yamanaka factors (Oct4, Sox2, Nanog) in mESCs and for MyoG and Pax7 in C2C12 cells, the reference was H3K4me2 CUT&Tag, using the DESeq2 results described above. For GATA2 in K562 cells, TF-CUT&Tag was compared directly to standard GATA2 CUT&Tag (control pA-Tn5). Peaks with a missing log2 fold change were removed, and peaks were filtered to canonical chromosomes. Peak sequences were extracted with BSgenome (v1.4.3) using the peak coordinates from the DESeq2 results tables (mm10 for mESC and C2C12, hg38 for K562). Peaks were ranked by DESeq2 log2 fold change and divided into bins containing equal numbers of peaks (monaLisa::bin, binmode = “equalN”). The number of peaks per bin was set for each comparison so that every heatmap displayed eight bins.

Motif enrichment in each bin was calculated with calcBinnedMotifEnrR, using position weight matrices and a minimum motif match score of 10. Each bin was tested against the peaks in all other bins, and the log2 enrichment and the −log10 adjusted p-value were calculated for each motif. Results were plotted with plotMotifHeatmaps, with the log2 enrichment color scale capped at 2 and the −log10 adjusted p-value scale capped at 10. Bins are ordered from lowest to highest log2 fold change.

Motifs were taken from the JASPAR 2024 (v0.99.7) vertebrate collection for all factors. For the Yamanaka factors, the motif set comprised of POU5F1, SOX2, NANOG, MYC/ MYCN, KLF4, and the POU5F1::SOX2 composite motif. For GATA2, the motif set comprised the GATA2, GATA1, GATA3 and GATA1::TAL1 motifs. For MyoG, myogenic E-box motifs Myf6, MyoD1 and MyoG were selected from JASPAR. For Pax7, motifs were selected from both JASPAR and HOCOMOCO v14.

### Classification of peaks as promoter-proximal or distal

Transcription start sites (TSSs) for mm10 were taken from a 1 bp TSS annotation and expanded by 1 kb on each side to give 2 kb windows centered on each TSS. Peaks were classified with BEDTools (via pybedtools) as promoter-proximal (“TSS”) if they overlapped at least 1 bp of any TSS window (bedtools intersect -u), and as “distal” otherwise. For Pax7, this classification was applied to the peaks favored by TF-CUT&Tag (K160A, 200 mM NaCl) and the peaks favored by standard CUT&Tag (control pA-Tn5, 300 mM NaCl) in the head-to-head signal comparisons at day 0 (Dn1) and day 1 (Dp1) of C2C12 differentiation, defined as described above. The percentage of each peak set falling in each class was plotted as stacked bar charts.

---

**Supplementary Figure 1.**
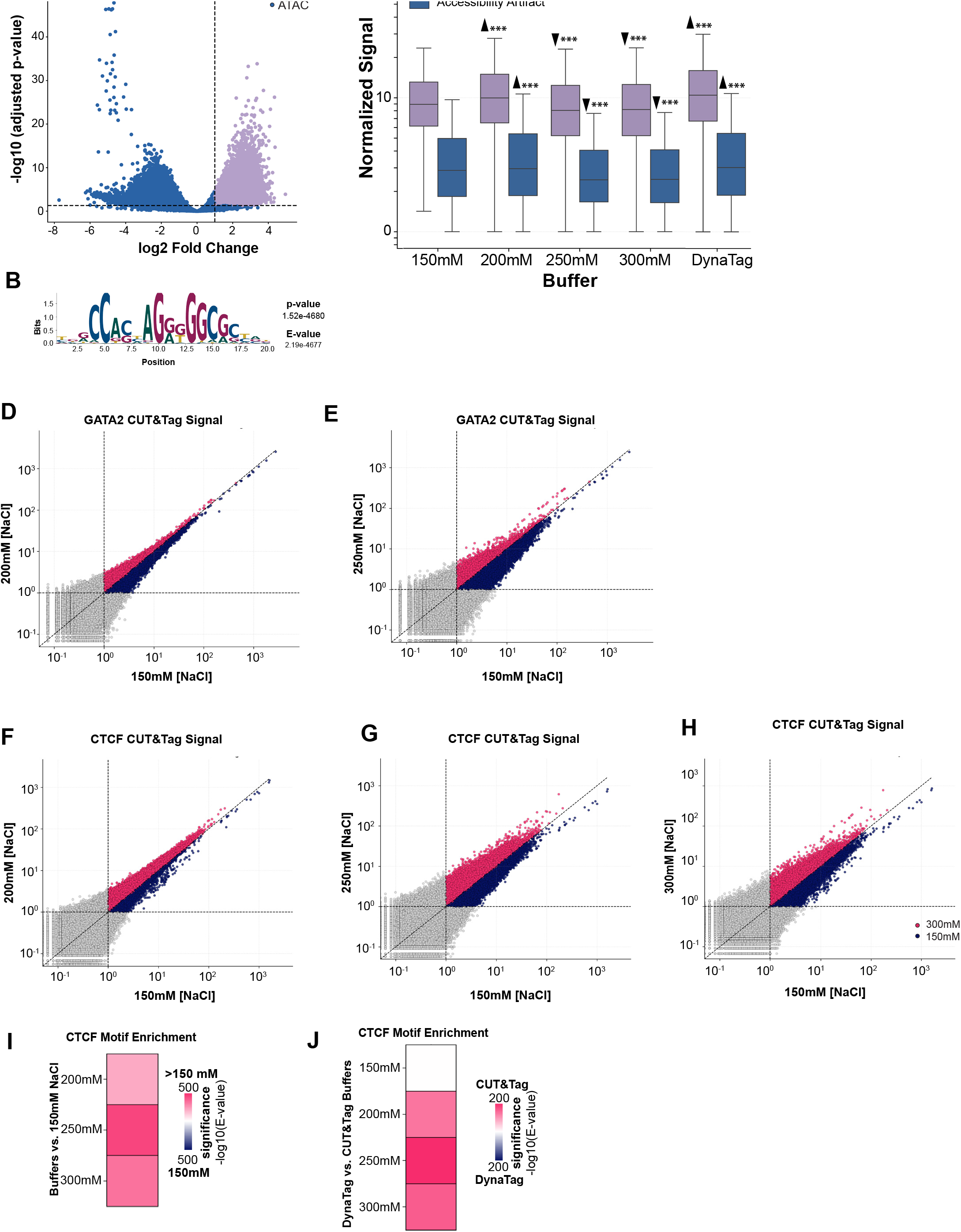
Analysis of CTCF and GATA2 CUT&Tag signal recovery and specificity across ionic conditions. **(A)** Volcano plot of DESeq2 results comparing CTCF CUT&Tag generated at 150 mM NaCl with the curated K562 ATAC-seq reference. Peaks with log2 fold change > 1 and adjusted P < 0.05 were classified as CTCF-associated peaks and are shown in purple; remaining peaks comprise the operational accessibility-associated reference set and are shown in blue. **(B)** Sequence logo and enrichment statistics from AME comparing CTCF-associated peaks with the accessibility-associated reference set defined in (A). **(C)** Boxplot of normalized CTCF CUT&Tag signal across CTCF-associated and accessibility-associated peak sets defined in (A). Statistical comparisons are between the 150 mM NaCl condition and the 200, 250, and 300 mM NaCl and DynaTag conditions, with arrowheads indicating the direction of significant differences. **(D–E)** Scatterplots comparing normalized signal across a merged GATA2 peak set between 150 mM NaCl and 200 mM (D) or 250 mM (E) NaCl. Each point represents one peak; peaks with normalized signal ≥1 on both axes are colored according to the direction of the signal difference. **(F–H)** Same as (D–E), but comparing normalized signal across a merged CTCF peak set between 150 mM NaCl and 200 mM (F), 250 mM (G), or 300 mM (H) NaCl. **(I–J)** Heatmaps showing differential enrichment of the CTCF motif by AME, represented as −log10(E-value). In (I), pink indicates enrichment among peaks favored by NaCl concentrations >150 mM and navy indicates enrichment among peaks favored at 150 mM NaCl. In (J), pink indicates enrichment among peaks favored by the indicated NaCl condition and navy indicates enrichment among peaks favored by DynaTag.

**Supplementary Figure 2.**
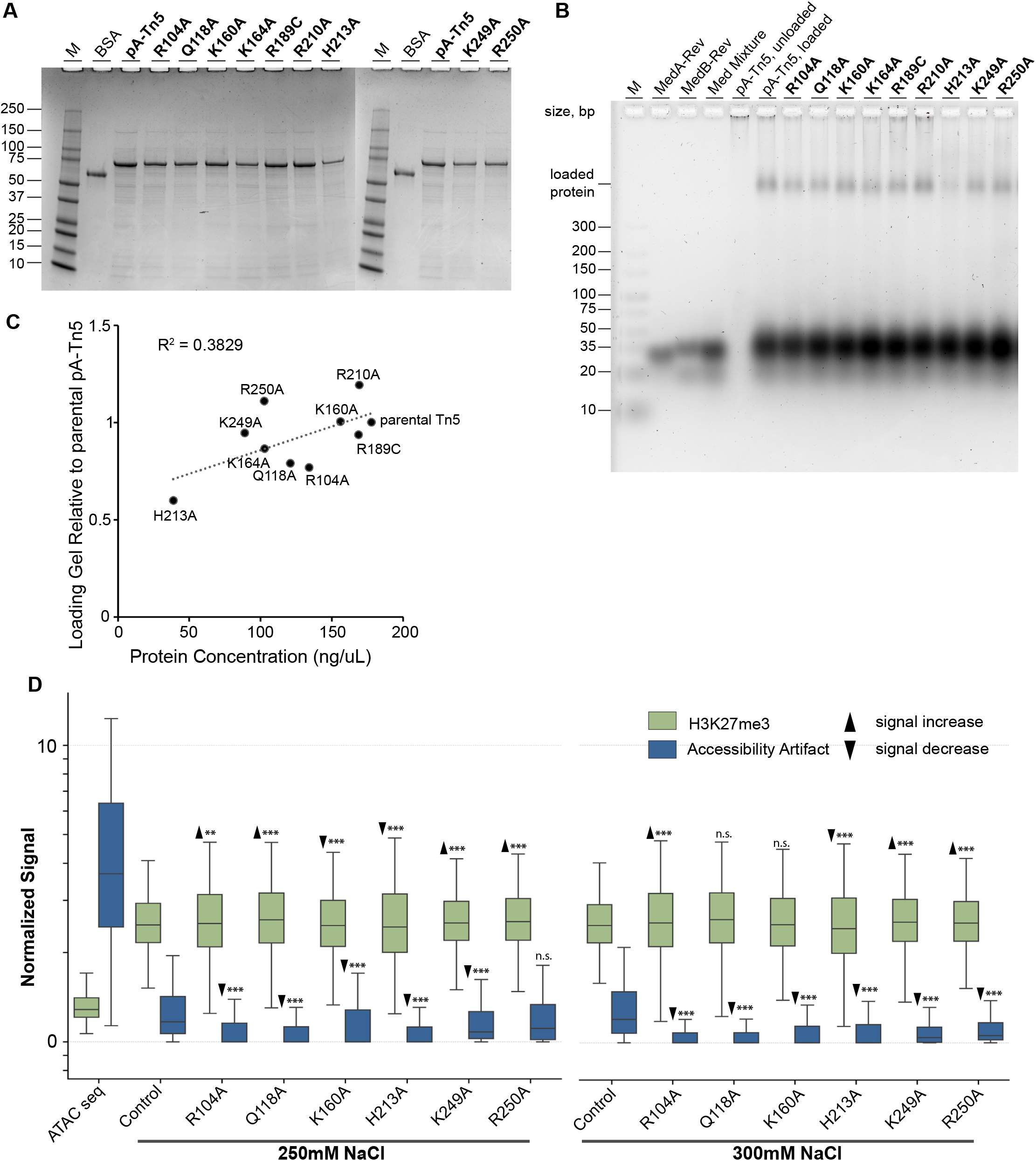
Generation and functional characterization of mutant pA-Tn5 variants. **(A)** SDS-PAGE analysis of purified parental and mutant pA-Tn5 preparations used to quantify protein abundance prior to oligonucleotide loading. **(B)** Gel-based analysis of oligonucleotide loading for parental and mutant pA-Tn5 variants, with the shifted species representing assembled pA-Tn5 transposomes. **(C)** Quantification of oligonucleotide loading efficiency for each pA-Tn5 variant, calculated by normalizing the amount of shifted oligonucleotide in (B) to the corresponding amount of purified pA-Tn5 protein in (A). **(D)** Boxplots of normalized H3K27me3-associated and accessibility-associated signal for parental and mutant pA-Tn5 variants under 250 and 300 mM NaCl conditions. Statistical comparisons are between parental and mutant pA-Tn5 variants, with arrowheads indicating the direction of significant differences.

**Supplementary Figure 3.**
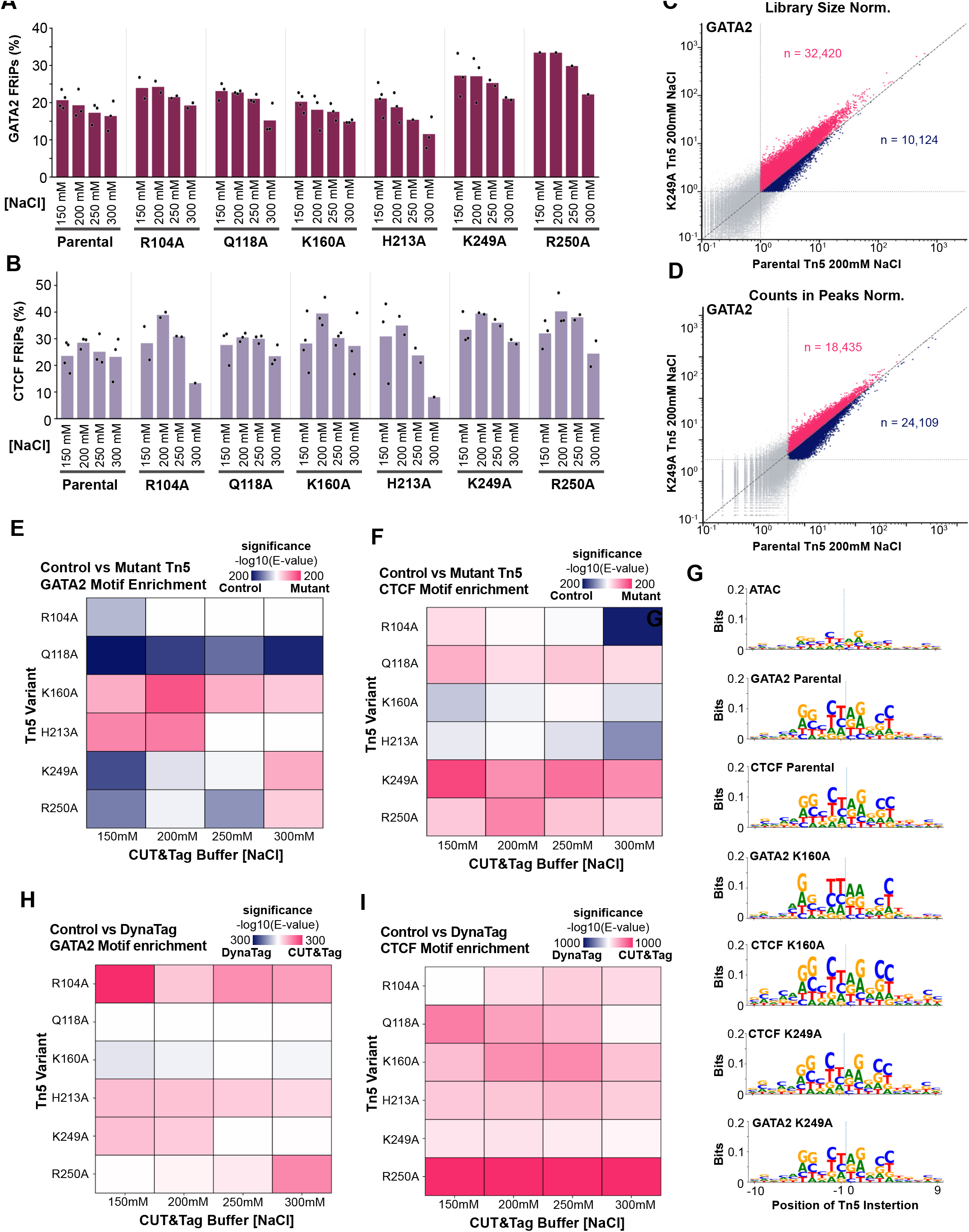
Library normalization and Tn5 insertion preferences do not explain TF-dependent variant performance. **(A,B)** Fraction of reads in peaks (FRiP) for GATA2 (A) and CTCF (B) CUT&Tag libraries generated with parental pA-Tn5 or the indicated variants across 150, 200, 250, and 300 mM NaCl. Bars summarize replicate FRiP values; black dots indicate individual libraries. **(C**,**D)** Scatterplots comparing GATA2 CUT&Tag signal for K249A and parental pA-Tn5 at 200 mM NaCl after normalization to the total eligible library fragments (C; library-size normalization) or to the fragments falling within the consensus peak set (D; counts-in-peaks normalization). Each point represents one peak; pink indicates K249A-favored peaks and navy indicates parental-enzyme-favored peaks among those passing the signal threshold. **(E**,**F)** AME heatmaps showing differential enrichment of GATA2 (E) and CTCF motifs between variant-favored and parental-enzyme-favored peak sets after normalization by fragments within the consensus peak set. Values are represented as signed −log10(E-value), with pink indicating enrichment in variant-favored peaks and blue indicating enrichment in parental-enzyme-favored peaks. The original whole-library-normalized signal threshold was retained to define peaks eligible for comparison. **(G)** Sequence logos showing Tn5 insertion-site sequence preferences in ATAC-seq and parental, K160A, and K249A CUT&Tag libraries for GATA2 and CTCF. Letter heights represent positional information in bits. **(H**,**I)** AME heatmaps comparing GATA2 (H) and CTCF (I) motif enrichment between NaCl-based CUT&Tag and DynaTag for the same indicated Tn5 variant. Pink indicates enrichment among NaCl-favored peaks and blue indicates enrichment among DynaTag-favored peaks, expressed as signed −log10(E-value).

**Supplementary Figure 4.**
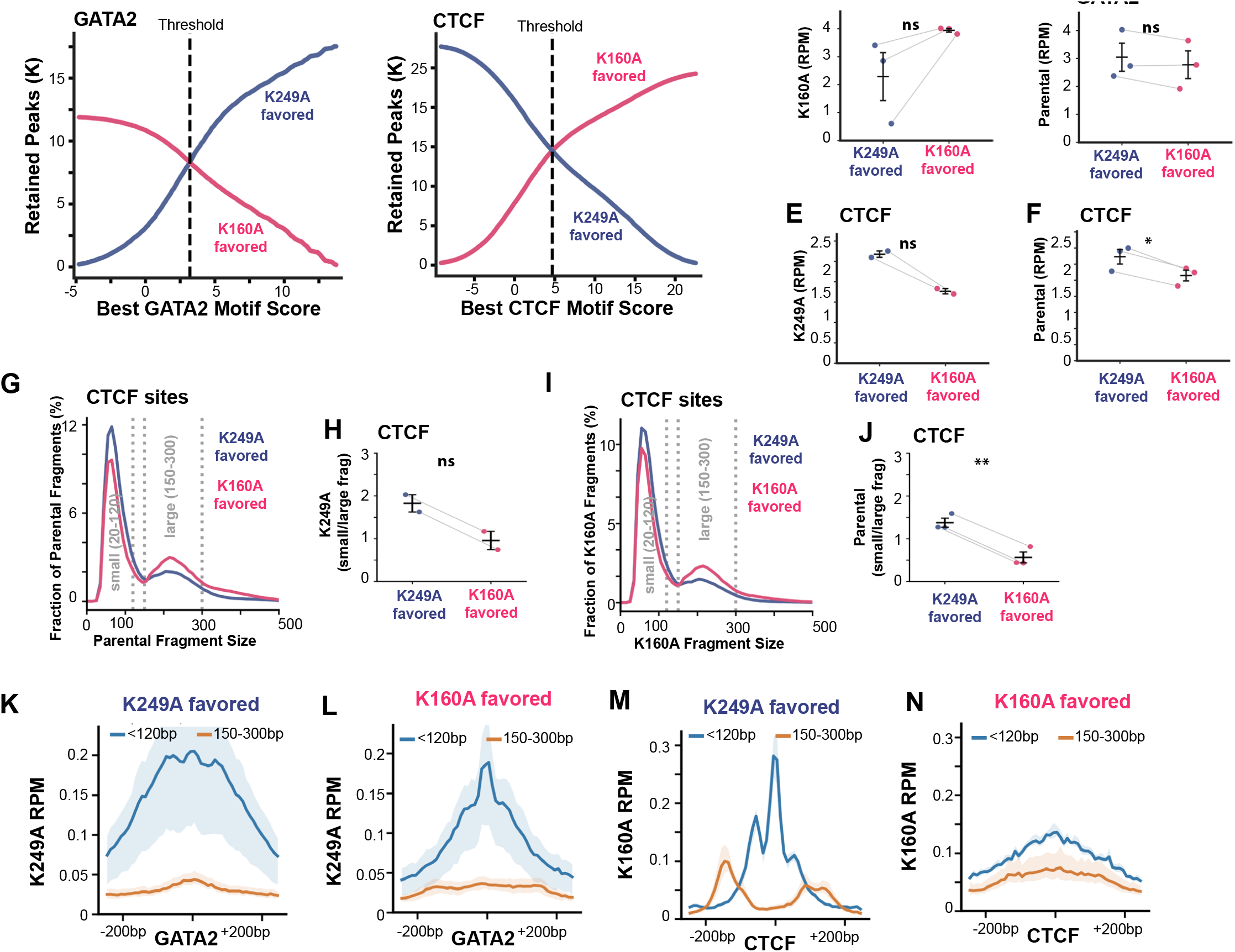
Signal recovery and fragment organization at K160A- and K249A-preferred sites. **(A,B)** Numbers of GATA2 (A) and CTCF (B) sites retained in the opposing variant-preference/motif-score populations as the minimum motif-score threshold is varied. Dashed vertical lines indicate the thresholds used to define the selected populations in Fig. 4A,B. **(C**,**D)** Paired-library comparisons of mean normalized CUT&Tag signal (RPM per peak) across K249A- and K160A-preferred GATA2 sites for K160A (C) and parental pA-Tn5 (D). **(E**,**F)** Corresponding comparisons across K249A- and K160A-preferred CTCF sites for K249A (E) and parental pA-Tn5 (F). In (C–F), each point represents one library; lines connect measurements from the same library and black symbols indicate mean ± SEM. **(G**,**H)** Fragment-length distributions (G) and paired-library ratios of short to nucleosome-sized fragments (H) from parental pA-Tn5 CUT&Tag at the selected CTCF sites. **(I**,**J)** Corresponding fragment-length distributions (I) and short-to-nu-cleosome-sized fragment ratios (J) from K160A CUT&Tag at the CTCF sites. In (G,I), blue denotes K249A-preferred sites and pink denotes K160A-preferred sites; dashed lines delimit short (20–119 bp) and nucleosome-sized (150–299 bp) fragment ranges. **(K**,**L)** Motif-centered mean K249A CUT&Tag fragment-midpoint profiles for K249A-preferred (K) and K160A-preferred (L) GATA2 sites. **(M**,**N)** Corresponding mean K160A CUT&Tag profiles for K249A-preferred (M) and K160A-preferred (N) CTCF sites. In (K–N), blue denotes short fragments and orange denotes nucleosome-sized fragments; shaded areas indicate the range among individual libraries.

**Supplementary Figure 5.**
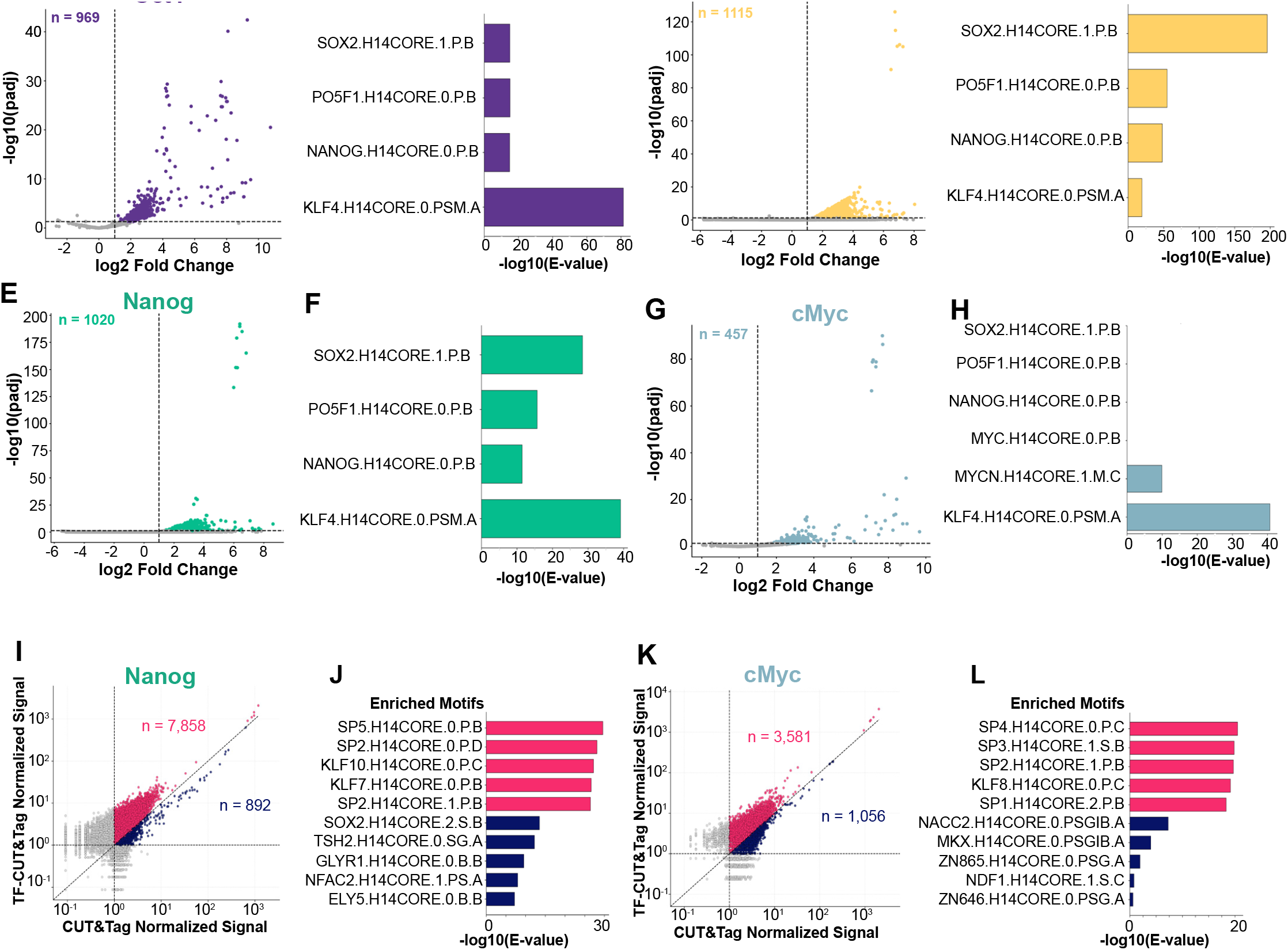
Pluripotency-associated motif enrichment in TF-CUT&Tag varies among transcription factors. **(A– H)** DESeq2 and motif enrichment analyses comparing TF-CUT&Tag with H3K4me2 CUT&Tag for Oct4 (A,B), Sox2 (C,D), Nanog (E,F), and cMyc (G,H). Volcano plots (A,C,E,G) show differential signal between TF-CUT&Tag and H3K4me2 CUT&Tag. Bar plots (B,D,F,H) show AME enrichment of the indicated pluripotency-associated motifs, represented as −log10(E-value). **(I**,**K)** Scatterplots comparing normalized signal across merged peak sets between TF-CUT&Tag and standard CUT&Tag for Nanog (I) and cMyc (K). Each point represents one peak. Peaks with normalized signal ≥1 on both axes were classified as TF-CUT&Tag-favored if they fell above the diagonal (y = x) or standard-CUT&Tag-favored if they fell below it, and are highlighted in pink and blue, respectively. **(J**,**L)** Bar plots showing differential motif enrichment by AME between TF-CUT&Tag-favored and standard-CUT&Tag-favored peak sets for Nanog (J) and cMyc (L), represented as −log10(E-value). Pink indicates motifs enriched among TF-CUT&Tag-favored peaks and blue indicates motifs enriched among standard-CUT&Tag-favored peaks.

## References

1 Lambert, S. A. et al. The Human Transcription Factors. Cell 172, 650–665 (2018). 10.1016/j.cell.2018.01.029

2 MacArthur, S. et al. Developmental roles of 21 Drosophila transcription factors are determined by quantitative differences in binding to an overlapping set of thousands of genomic regions. Genome Biol 10, R80 (2009). 10.1186/gb-2009-10-7-r80

3 Zaret, K. S. & Carroll, J. S. Pioneer transcription factors: establishing competence for gene expression. Genes Dev 25, 2227–2241 (2011). 10.1101/gad.176826.111

4 Mayran, A. et al. Pioneer and nonpioneer factor cooperation drives lineage specific chromatin opening. Nat Commun 10, 3807 (2019). 10.1038/s41467-019-11791-9

5 Kribelbauer, J. F., Rastogi, C., Bussemaker, H.J. & Mann, R. S. Low-Affinity Binding Sites and the Transcription Factor Specificity Paradox in Eukaryotes. Annu Rev Cell Dev Biol 35, 357–379 (2019). 10.1146/annurev-cellbio-100617-062719

6 Li, X. Y. et al. The role of chromatin accessibility in directing the widespread, overlapping patterns of Drosophila transcription factor binding. Genome Biol 12, R34 (2011). 10.1186/gb-2011-12-4-r34

7 Abidi, A. A. et al. Unstructured transcription factor interactions enable emergent specificity. Science 392, eaeb6487 (2026). 10.1126/science.aeb6487

8 Crocker, J., Noon, E. P. & Stern, D. L. The Soft Touch: Low-Affinity Transcription Factor Binding Sites in Development and Evolution. Curr Top Dev Biol 117, 455–469 (2016). 10.1016/bs.ctdb.2015.11.018

9 Fan, V. B., Abidi, A. A., Graham, T. G. W., Darzacq,X. & Staller, M. V. Short activation domains control chromatin association of transcription factors. Elife 15 (2026). 10.7554/eLife.105776

10 Teytelman, L., Thurtle, D. M., Rine, J. & van Oudenaarden, A. Highly expressed loci are vulnerable to misleading ChIP localization of multiple unrelated proteins. Proc Natl Acad Sci U S A 110, 18602–18607 (2013). 10.1073/pnas.1316064110

11 Liu, N. et al. Direct Promoter Repression by BCL11AControls the Fetal to Adult Hemoglobin Switch. Cell 173, 430–442 e417 (2018). 10.1016/j.cell.2018.03.016

12 Ahmad, K. et al. Enzyme tethering for in situ epigenomics. Nature Reviews Methods Primers 6, 42 (2026). 10.1038/s43586-026-00491-6

13 Skene, P. J. & Henikoff, S. An efficient targeted nuclease strategy for high-resolution mapping of DNA binding sites. Elife 6 (2017). 10.7554/eLife.21856

14 Hainer, S. J., Boskovic, A., McCannell, K. N., Rando, O. J. & Fazzio, T. G. Profiling of Pluripotency Factors in Single Cells and Early Embryos. Cell 177, 1319–1329 e1311 (2019). 10.1016/j.cell.2019.03.014

15 Chi, W. Y. et al. Single-cell mapping of regulatory DNA-protein interactions. Cell 189, 3801–3816 e3811 (2026). 10.1016/j.cell.2026.05.014

16 Kaya-Okur, H. S., Janssens, D. H., Henikoff, J. G., Ahmad, K. & Henikoff, S. Efficient low-cost chromatin profiling with CUT&Tag. Nat Protoc 15, 3264–3283 (2020). 10.1038/s41596-020-0373-x

17 Kaya-Okur, H. S. et al. CUT&Tag for efficient epigenomic profiling of small samples and single cells. Nat Commun 10, 1930 (2019). 10.1038/s41467-019-09982-5

18 Janssens, D. H. et al. Automated CUT&Tag profiling of chromatin heterogeneity in mixed-lineage leukemia. Nat Genet 53, 1586–1596 (2021). 10.1038/s41588-021-00941-9

19 Wu, S. J. et al. Single-cell CUT&Tag analysis of chromatin modifications in differentiation and tumor progression. Nat Biotechnol 39, 819–824 (2021). 10.1038/s41587-021-00865-z

20 Bartosovic, M., Kabbe, M. & Castelo-Branco, G. Single-cell CUT&Tag profiles histone modifications and transcription factors in complex tissues. Nat Biotechnol 39, 825–835 (2021). 10.1038/s41587-021-00869-9

21 Meers, M. P., Llagas, G., Janssens, D. H., Codomo,C. A. & Henikoff, S. Multifactorial profiling of epigenetic landscapes at single-cell resolution using MulTI-Tag. Nat Biotechnol 41, 708–716 (2023). 10.1038/s41587-022-01522-9

22 Stuart, T. et al. Nanobody-tethered transposition enables multifactorial chromatin profiling at single-cell resolution. Nat Biotechnol 41, 806–812 (2023). 10.1038/s41587-022-01588-5

23 Barcenas-Walls, J. R. et al. Nano-CUT&Tag for multimodal chromatin profiling at single-cell resolution. Nat Protoc 19, 791–830 (2024). 10.1038/s41596-023-00932-6

24 Bartosovic, M. & Castelo-Branco, G. Multimodal chromatin profiling using nanobody-based single-cell CUT&Tag. Nat Biotechnol 41, 794–805 (2023). 10.1038/s41587-022-01535-4

25 Liu, M. et al. Genome-coverage single-cell histone modifications for embryo lineage tracing. Nature 640, 828–839 (2025). 10.1038/s41586-025-08656-1

26 Hunold, P. et al. DynaTag for efficient mapping of transcription factors in low-input samples and at single-cell resolution. Nat Commun 16, 6585 (2025). 10.1038/s41467-025-61797-9

27 Gradman, R. J., Ptacin, J. L., Bhasin, A., Reznikoff,W. S. & Goryshin, I. Y. A bifunctional DNA binding region in Tn5 transposase. Mol Microbiol 67, 528–540 (2008). 10.1111/j.1365-2958.2007.06056.x

28 Corces, M. R. et al. An improved ATAC-seq protocol reduces background and enables interrogation of frozen tissues. Nat Methods 14, 959–962 (2017). 10.1038/nmeth.4396

29 Krohannon, A., Srivastava, M., Sangani, N. & Janga,S. C. Regulation Ratio: A Singular Multi-Omic Measurement of Gene Regulatory Mechanisms. Comput Struct Biotechnol J 35, 0044 (2026). 10.34133/csbj.0044

30 Consortium, E. P. An integrated encyclopedia of DNA elements in the human genome. Nature 489, 57–74 (2012). 10.1038/nature11247

31 Andreeva, T. V. et al. Na(+) and K(+) Ions Differently Affect Nucleosome Structure, Stability, and Interactions with Proteins. Microsc Microanal 28, 243–253 (2022). 10.1017/S1431927621013751

32 Chen, C. W. et al. DOT1L inhibits SIRT1-mediated epigenetic silencing to maintain leukemic gene expression in MLL-rearranged leukemia. Nat Med 21, 335–343 (2015). 10.1038/nm.3832

33 Henikoff, S., Henikoff, J. G., Kaya-Okur, H. S. & Ahmad, K. Efficient chromatin accessibility mapping in situ by nucleosome-tethered tagmentation. Elife 9 (2020). 10.7554/eLife.63274

34 Zammit, P. S. et al. Pax7 and myogenic progression in skeletal muscle satellite cells. J Cell Sci 119, 1824–1832 (2006). 10.1242/jcs.02908

35 Hargreaves, V. V. & Schleif, R. F. The salt dependence of the interferon regulatory factor 1 DNA binding domain binding to DNA reveals ions are localized around protein and DNA. Biochemistry 47, 4119–4128 (2008). 10.1021/bi702082q

36 Sakong, S., Fierz, B. & Suter, D. M. Electrostatic properties of disordered regions control transcription factor search and pioneer activity. Nat Commun 17 (2026). 10.1038/s41467-026-69284-5

37 Chen, J. et al. Single-molecule dynamics of enhanceosome assembly in embryonic stem cells. Cell 156, 1274–1285 (2014). 10.1016/j.cell.2014.01.062

38 Hansen, A. S., Pustova, I., Cattoglio, C., Tjian, R. & Darzacq, X. CTCF and cohesin regulate chromatin loop stability with distinct dynamics. Elife 6 (2017). 10.7554/eLife.25776

39 Pelletier, A. et al. Pax7 pioneer factor action requires both paired and homeo DNA binding domains.Nucleic Acids Res 49, 7424–7436 (2021). 10.1093/nar/gkab561

40 Yoshida, N., Yoshida, S., Koishi, K., Masuda, K. & Nabeshima, Y. Cell heterogeneity upon myogenic differentiation: down-regulation of MyoD and Myf-5 generates ‘reserve cells’. J Cell Sci 111 (Pt 6), 769–779 (1998). 10.1242/jcs.111.6.769

41 Olguin, H. C. & Olwin, B. B. Pax-7 up-regulation inhibits myogenesis and cell cycle progression in satellite cells: a potential mechanism for self-renewal. Dev Biol 275, 375–388 (2004). 10.1016/j.ydbio.2004.08.015

42 Hennig, B. P. et al. Large-Scale Low-Cost NGS Library Preparation Using a Robust Tn5 Purification and Tagmentation Protocol. G3 (Bethesda) 8, 79–89 (2018). 10.1534/g3.117.300257

43 Nanda, A. S. et al. Direct transposition of native DNA for sensitive multimodal single-molecule sequencing. Nat Genet 56, 1300–1309 (2024). 10.1038/s41588-024-01748-0

44 Sun, W., Baird, E., Ashbaugh, H. J., Eiken, A.P. & Janssens, D. H. CoCUT&Tag maps linked chromatin states at single-molecule, single-cell resolution. bioRxiv (2026). 10.64898/2026.04.27.721191

